## Supplementary Information for "Genetics and evidence for balancing selection of a sex-linked colour polymorphism in a songbird"

Kim *et al*.

This file contains:

- Supplementary Figures 1-20
- Supplementary Tables 1-14
- Supplementary Methods
- Supplementary References

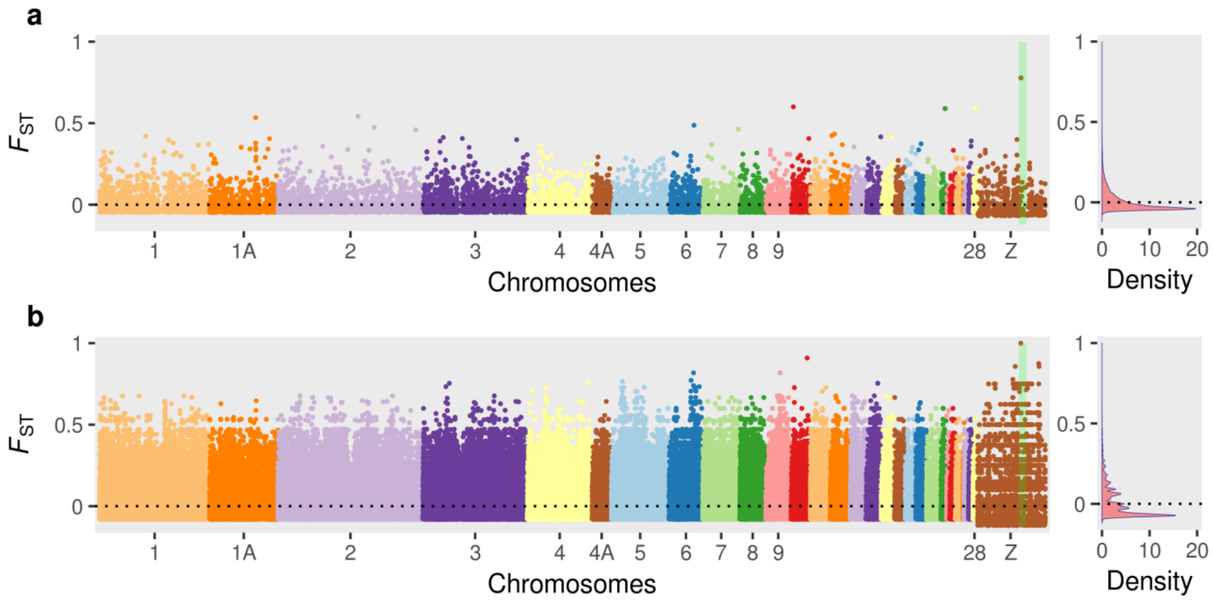


**Supplementary Figure 1 |** Pattern of genomic differentiation between black and red Gouldian finches. We treated each morph as a separate population. A value for each SNP is plotted against its position in the zebra finch genome assembly. The region shaded in green on the Z chromosome represents the candidate interval (~7.2 cM) for the *Red* locus based on a previous linkage mapping study[1]. SNPs with < 80% genotypes per morph or an unmapped position were excluded. Right panels show the frequency distribution of the $F_{ST}$ values. **a**, Site-by-site $F_{ST}$ in the wild population ($n_{\text{black male}}$ = 22, $n_{\text{red female}}$ = 8 and $n_{\text{red male}}$ = 2) using 14,800 SNPs (14,289 autosomal and 511 Z-linked) from RADSeq. Two SNPs (Z: 46,561,597–46,561,598) within the candidate interval are mapped to the *M1* region (Fig. 1, Supplementary Table2). **b**, Site-by-site $F_{ST}$ in the captive population ($n_{\text{black male}}$ = 4, $n_{\text{black female}}$ = 3, $n_{\text{red female}}$ = 5 and $n_{\text{yellow female}}$ = 5) using 530,163 SNPs (519,673 autosomal and 10,490 Z-linked) from WGS. A single SNP (Z: 46,580,103) within the candidate interval is mapped close to *M3* (Fig. 1, Supplementary Table2). The composite estimates of $F_{ST}$ using all SNPs were $F_{ST}$ = 0.006 and $F_{ST}$ = 0.017 for the wild and the captive populations, respectively. Note that, as female birds have hemizygous sex chromosome, smaller numbers of alleles were included in the analyses of the Z chromosome (wild population: 44 and 12 alleles for black and red birds respectively; captive population: 11 and 10 alleles for black and red birds respectively) than in the autosomal chromosomes.

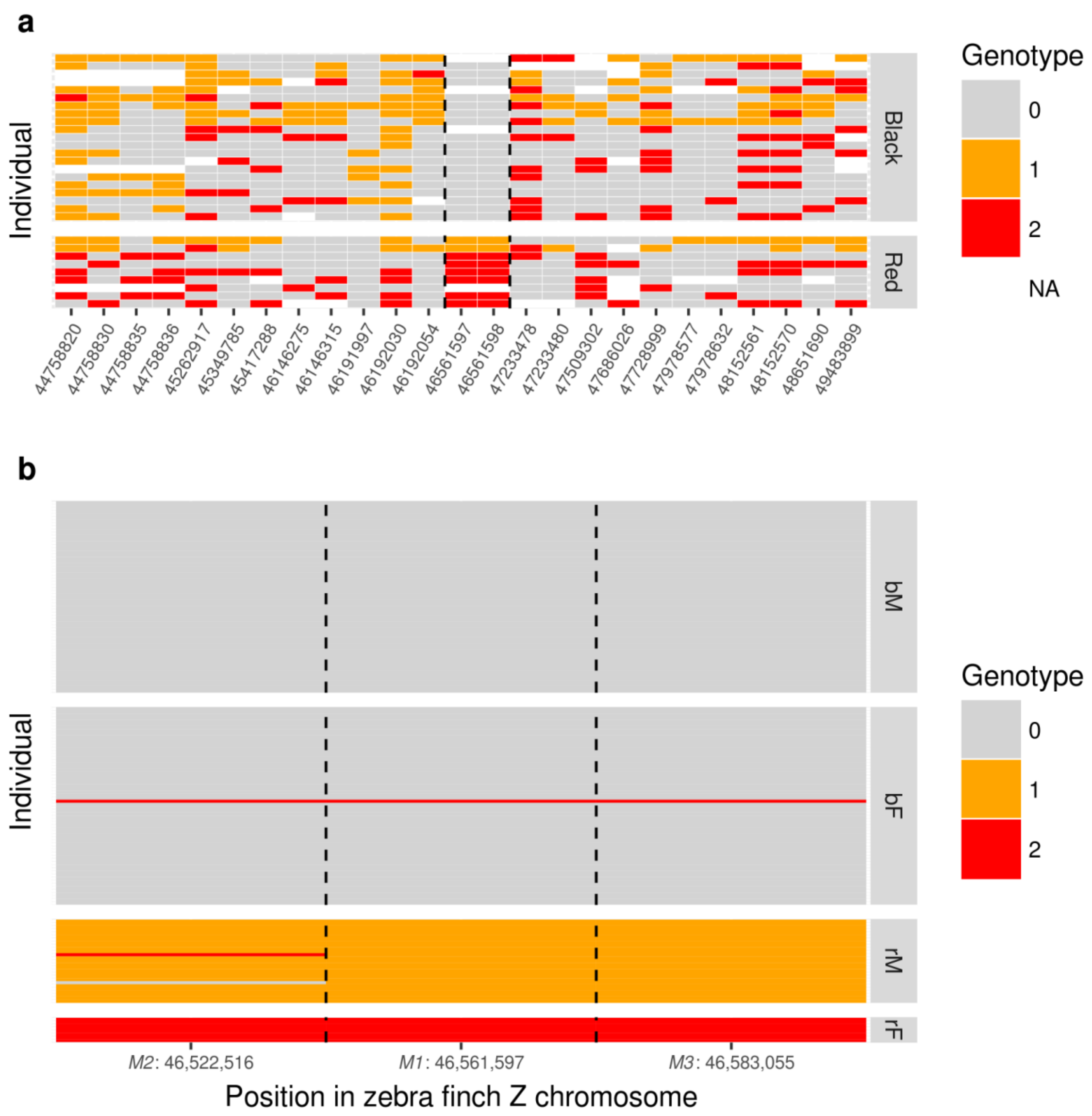


**Supplementary Figure 2 |** Genotypes of Gouldian finches from a wild population. Positions are based on the Z chromosome of the zebra finch. Major alleles in the samples were coded as '0' representing homozygotes for reference allele, '2' representing homozygotes for the alternative allele and '1' being a heterozygote. Females were coded as one of the homozygotes. **a**, RADSeq genotypes of 30 Gouldian finches (black, $n_{\text{male}}$ = 21; red, $n_{\text{male}}$ = 2, $n_{\text{female}}$ = 7; two individuals with a high level of missing data were excluded) around the *Red* locus (see Fig. 1). The two consecutive SNPs in *M1* (46,561,597–46,561,598 bp) are located in the *Red* locus and were used to genotype additional samples using an allele-specific PCR. **b**, Genotypes of 161 Gouldian finches (black male, $n_{\text{bM}}$ = 62; black female, $n_{\text{bF}}$ = 64; red male, $n_{\text{rM}}$ = 27; red female, $n_{\text{rF}}$ = 8) for the three markers (*M1*–*M3*) across the *Red* locus (see Supplementary Table 2). Two recombination events between black- and red-linked alleles are visible in the genotypes of red males. A single black female with red-linked alleles is likely to represent a phenotyping error due to the less bright head colours in females in the wild population.

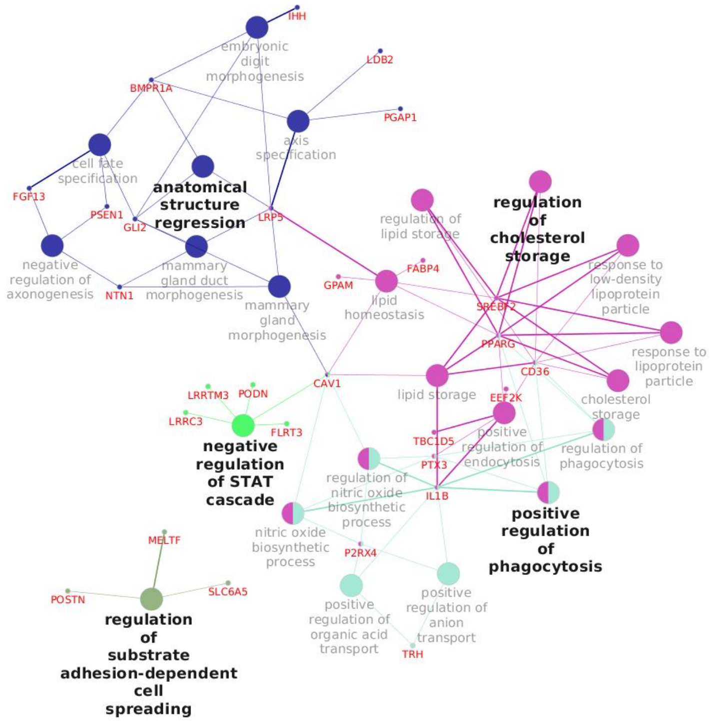


**Supplementary Figure 3 |** A functionally-grouped annotation network of genes differentially-expressed in the regenerating feather follicles of black ($n_{\text{male}}$ = 3) and red ($n_{\text{male}}$ = 2, $n_{\text{female}}$ = 1) birds. Five groups of GO terms are represented by circles of different colour. The group leading terms are shown in bold black text and the genes connecting the terms are shown in red. Most notably, one of the major GO groups was involved in cell fate/morphology specification. In particular, *BMP receptor type 1A* (*BMPR1A*) and *fibroblast growth factor 13* (*FGF-13*) in this group are likely to regulate feather/hair development[2-5]. Another major GO group is involved in lipid storage that may act in lipid-rich keratinocytes that are characteristic of birds[6]. Intriguingly, the genes that connect GO terms in this group (e.g. *PPARγ*, *CD36*, *IL1B*) appear to be regulated by interaction with astaxanthin[7,8], a red carotenoid found in red plumage of the Gouldian finch[9], suggesting that differentially accumulated pigments might also be important in the development of different types of feather. In addition, *CYP2J19* (previously annotated as *CYP2J2-1* in zebra finch) was highly expressed in red birds, suggesting that the astaxanthin may be produced by ketolase conversion in the red morph[10]. See Supplementary Data 1 for the full list of differentially expressed genes.

```
                  M   L   N   Q   R   I   H   P   G   M   L   L   I   L   M   F   L   C   H   F
Zebra finch ATG TTA AAT CAG AGA ATC CAC CCG GGC ATG CTC TTA ATC CTG ATG TTT CTG TGC CAC TTC  [  60]
Black F102  ... ... ... ... ... ... ... ... ... ... ... ... ... ... ... ... ... ... ... ...  [  60]
Red   F125  ... ... ... ... ... ... ... ... ... ... ... ... ... ... ... ... ... ... ... ...  [  60]
                  M   E   D   H   T   V   Q   A   G   N   C   W   L   R   Q   A   R   N   G   R
Zebra finch ATG GAA GAT CAC ACA GTG CAG GCT GGG AAC TGC TGG CTC CGC CAG GCG CGG AAC GGG CGC  [ 120]
Black F102  ... ... ... ... ... ... ... ... ... ... ... ... ... ..G ... ... ... ... ... ...  [ 120]
Red   F125  ... ... ... ... ... ... ... ... ... ... ... ... ... ..G ... ... ... ... ... ...  [ 120]
                  C   Q   V   L   Y   K   T   D   L   S   K   E   E   C   C   K   S   G   R   L
Zebra finch TGC CAG GTC CTC TAC AAA ACC GAC CTC AGC AAG GAG GAG TGC TGC AAG AGC GGC CGC CTG  [ 180]
Black F102  ... ... ... ... ... ... ... ... ... ... ... ... ... ... ... ... ... ... ... ...  [ 180]
Red   F125  ... ... ... ... ... ... ... ... ... ... ... ... ... ... ... ... ... ... ... ...  [ 180]
                  T   T   S   W   T   A   E   D   V   N   D   N   T   L   F   K   W   M   I   F
Zebra finch ACC ACG TCG TGG ACG GCG GAG GAC GTC AAC GAC AAT ACG CTT TTT AAG TGG ATG ATT TTT  [ 240]
Black F102  ... ... ... ... ... ... ... ... ... ... ... ... ... ... ... ... ... ... ... ...  [ 240]
Red   F125  ... ... ... ... ... ... ... ... ... ... ... ... ... ... ... ... ... ... ... ...  [ 240]
                  N   G   G   A   P   N   C   I   P   C   K   E   T   C   E   N   V   D   C   G
Zebra finch AAT GGG GGA GCC CCA AAC TGC ATC CCA TGC AAA GAA ACA TGC GAG AAT GTG GAC TGT GGA  [ 300]
Black F102  ... ... ... ... ... ... ... ... ... ... ... ... ... ... ... ... ... ... ... ...  [ 300]
Red   F125  ... ... ... ... ... ... ... ... ... ... ... ... ... ... ... ... ... ... ... ...  [ 300]
                  P   G   K   K   C   K   M   N   K   K   N   K   P   R   C   V   C   A   P   D
Zebra finch CCC GGG AAG AAA TGT AAA ATG AAC AAG AAG AAC AAA CCT CGG TGT GTT TGT GCT CCG GAT  [ 360]
Black F102  ... ... ... ... ... ... ... ... ... ... ... ... ... ... ... ... ... ... ... ...  [ 360]
Red   F125  ... ... ... ... ... ... ... ... ... ... ... ... ... ... ... ... ... ... ... ...  [ 360]
                  C   S   N   I   T   W   K   G   P   V   C   G   L   D   G   K   T   Y   R   N
Zebra finch TGC TCT AAT ATC ACT TGG AAG GGC CCC GTG TGT GGC TTA GAT GGG AAA ACC TAC AGG AAC  [ 420]
Black F102  ... ... ... ... ... ... ... ..T ... ... ... ... ... ... ... ... ... ... ... ...  [ 420]
Red   F125  ... ... ... ... ... ... ... ..T ... ... ... ... ... ... ... ... ... ... ... ...  [ 420]
                  E   C   A   L   L   K   A   R   C   K   E   Q   P   E   L   E   V   Q   Y   Q
Zebra finch GAG TGC GCC CTT CTC AAA GCC AGA TGT AAA GAA CAG CCT GAA CTT GAA GTC CAG TAT CAG  [ 480]
Black F102  ... ... ... ... ... ... ... ... ... ... ... ... ... ... ... ... ... ... ... ...  [ 480]
Red   F125  ... ... ... ... ... ... ... ... ... ... ... ... ... ... ... ... ... ... ... ...  [ 480]
                  G   K   C   K   K   T   C   R   D   V   L   C   P   G   S   S   T   C   V   V
Zebra finch GGC AAA TGC AAA AAA ACC TGC AGA GAT GTC TTA TGC CCA GGC AGC TCC ACA TGT GTG GTT  [ 540]
Black F102  ... ... ... ... ... ... ... ... ... ... ... ... ... ... ... ... ... ... ... ...  [ 540]
Red   F125  ... ... ... ... ... ... ... ... ... ... ... ... ... ... ... ... ... ... ... ...  [ 540]
                  D   Q   T   N   N   A   Y   C   V   T   C   N   R   I   C   P   E   P   T   S
Zebra finch GAC CAA ACC AAC AAT GCA TAC TGC GTG ACA TGT AAC CGA ATT TGT CCA GAG CCT ACC TCC  [ 600]
Black F102  ... ... ... ... ... ..C ... ..T ... ... ... ... ... ... ... ... ... ... ... ...  [ 600]
Red   F125  ... ... ... ... ... ..C ... ..T ... ... ... ... ... ... ... ... ... ... ... ...  [ 600]
                  A   E   Q   Y   L   C   G   N   D   G   I   T   Y   A   S   A   C   H   L   R
Zebra finch GCT GAA CAG TAT CTC TGT GGG AAT GAC GGC ATA ACT TAC GCC AGT GCC TGC CAC CTG AGG  [ 660]
Black F102  ... ..G ... ... ... ... ... ... ... ... ... ... ... ... ... ... ... ... ... ...  [ 660]
Red   F125  ... ..G ... ... ... ... ... ... ... ... ... ... ... ... ... ... ... ... ... ...  [ 660]
                  K   A   T   C   L   L   G   R   S   I   G   L   A   Y   E   G   K   C   V   K
Zebra finch AAA GCT ACC TGC CTC CTG GGA AGA TCC ATT GGA TTA GCC TAT GAA GGA AAA TGT GTC AAA  [ 720]
Black F102  ... ... ... ... ... ... ... ... ... ... ... ... ... ... ... ... ... ..C ... ...  [ 720]
Red   F125  ... ... ... ... ... ... ... ... ... ... ... ... ... ... ... ... ... ..C ... ...  [ 720]
                  A   K   S   C   E   D   I   Q   C   S   A   G   K   K   C   L   W   D   F   K
Zebra finch GCC AAA TCC TGT GAA GAC ATT CAA TGC AGT GCT GGG AAG AAA TGC TTG TGG GAT TTT AAG  [ 780]
Black F102  ... ... ... ... ... ... ... ... ... ... ... ... ... ... ... ... ... ... ... ...  [ 780]
Red   F125  ... ... ... ... ... ... ... ... ... ... ... ... ... ... ... ... ... ... ... ...  [ 780]
                  V   G   R   G   R   C   A   L   C   D   E   L   C   P   E   S   K   S   E   E
Zebra finch GTT GGC AGA GGT CGG TGT GCC CTC TGC GAT GAG CTA TGC CCT GAA AGC AAG TCA GAA GAA  [ 840]
Black F102  ... ... ... ... ... ... ... ... ..T ... ... ... ... ... ... ... ... ... ... ...  [ 840]
Red   F125  ... ... ... ... ... ... ... ... ..T ... ... ... ... ... ... ... ... ... ... ...  [ 840]
                  A   V   C   A   S   D   N   T   T   Y   P   S   E   C   A   M   K   E   A   A
Zebra finch GCA GTG TGT GCC AGC GAT AAC ACA ACT TAC CCA AGC GAG TGT GCC ATG AAA GAG GCA GCT  [ 900]
Black F102  ... ... ... ... ... ... ... ... ... ... ... ... ... ... ... ... ... ... ... ...  [ 900]
Red   F125  ... ... ... ... ... ... ... ... ... ... ... ... ... ... ... ... ... ... ... ...  [ 900]
                  C   S   M   G   V   L   L   E   V   K   H   S   G   S   C   N   S   I   N   E
Zebra finch TGC TCC ATG GGT GTG CTT CTA GAA GTT AAG CAC TCT GGA TCT TGC AAC TCA ATT AAT GAA  [ 960]
Black F102  ... ... ... ... ... ... ... ... ... ... ... ... ... ... ... ... ... ... ... ...  [ 960]
Red   F125  ... ... ... ... ... ... ... ... ... ... ... ... ... ... ... ... ... ... ... ...  [ 960]
                  D   P   E   D   E   E   E   D   E   D   Q   D   Y   S   F   P   I   S   S   I
Zebra finch GAC CCA GAA GAT GAA GAG GAA GAT GAA GAC CAG GAC TAC AGC TTT CCT ATA TCT TCC ATT  [1020]
Black F102  ... ... ... ... ... ... ... ... ... ... ... ... ... ... ... ... ... ... ... ...  [1020]
Red   F125  ... ... ... ... ... ... ... ... ... ... ... ... ... ... ... ... ... ... ... ...  [1020]
                  L   E   W   *
Zebra finch CTA GAG TGG TAA  [1032]
Black F102  ... ... ... ...  [1032]
Red   F125  ... ... ... ...  [1032]
```

**Supplementary Figure 4 |** Coding sequences of *follistatin* (*FST*) in 16 female Gouldian finches ($n_{\mathrm{black}}$ = 8, $n_{\mathrm{red}}$ = 8) compared to the zebra finch reference sequence. Only two representative individuals are shown because there was no polymorphism among Gouldian finches. Six exonic sequences of zebra finch were obtained from Ensembl (https://www.ensembl.org/). Amino acid sequences are shown at the top of the alignment. Identical sequences are indicated by dots and seven synonymous substitutions between zebra finch and Gouldian finch are shown in bold characters. The position of the last sequence in each line is shown in brackets on the right. Sequences were obtained from hemizygous Z chromosomes in females.

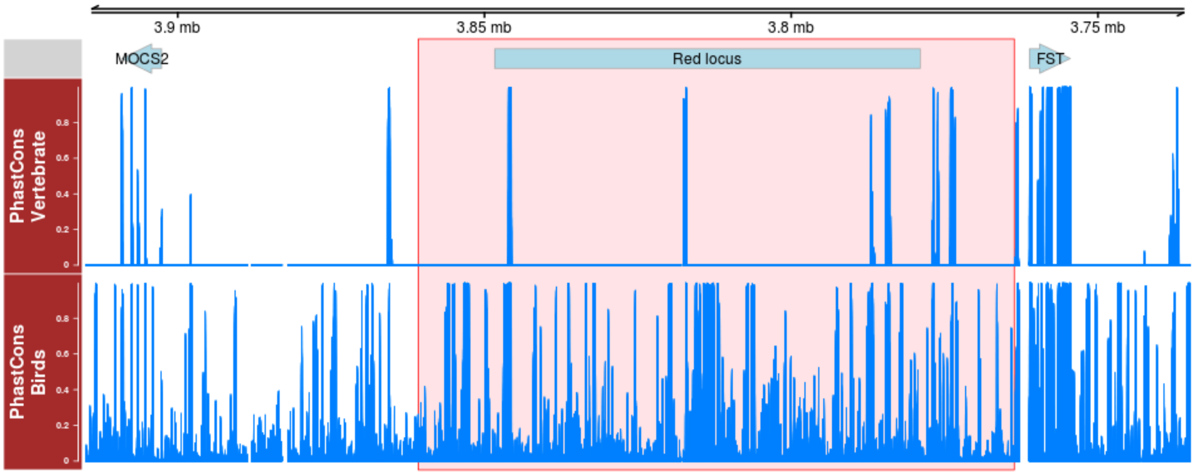


**Supplementary Figure 5 |** Evolutionarily conserved regions between *MOCS2* and *FST*. In the top panel, genomic positions of the *Red* locus and two flanking genes are displayed based on the genome assembly of the medium ground finch (*Geospiza fortis: geoFor1*, JH739925: 3,735,000–3,915,000 bp). Note that the reverse-oriented assembly is shown to allow comparison with the zebra finch genome assembly, as in Fig. 1-3. PhastCons values obtained from the UCSC Genome Browser Gateway (see Methods) based on the alignments of three vertebrates including medium ground finch, human and mouse (middle panel), and five birds including medium ground finch, zebra finch, budgerigar, chicken and turkey (bottom panel), are shown between the transcripts of *MOCS2* and *FST* (left and right sides of the top panel, respectively). The position of the 100-kbp sequenced region is highlighted in a red box.

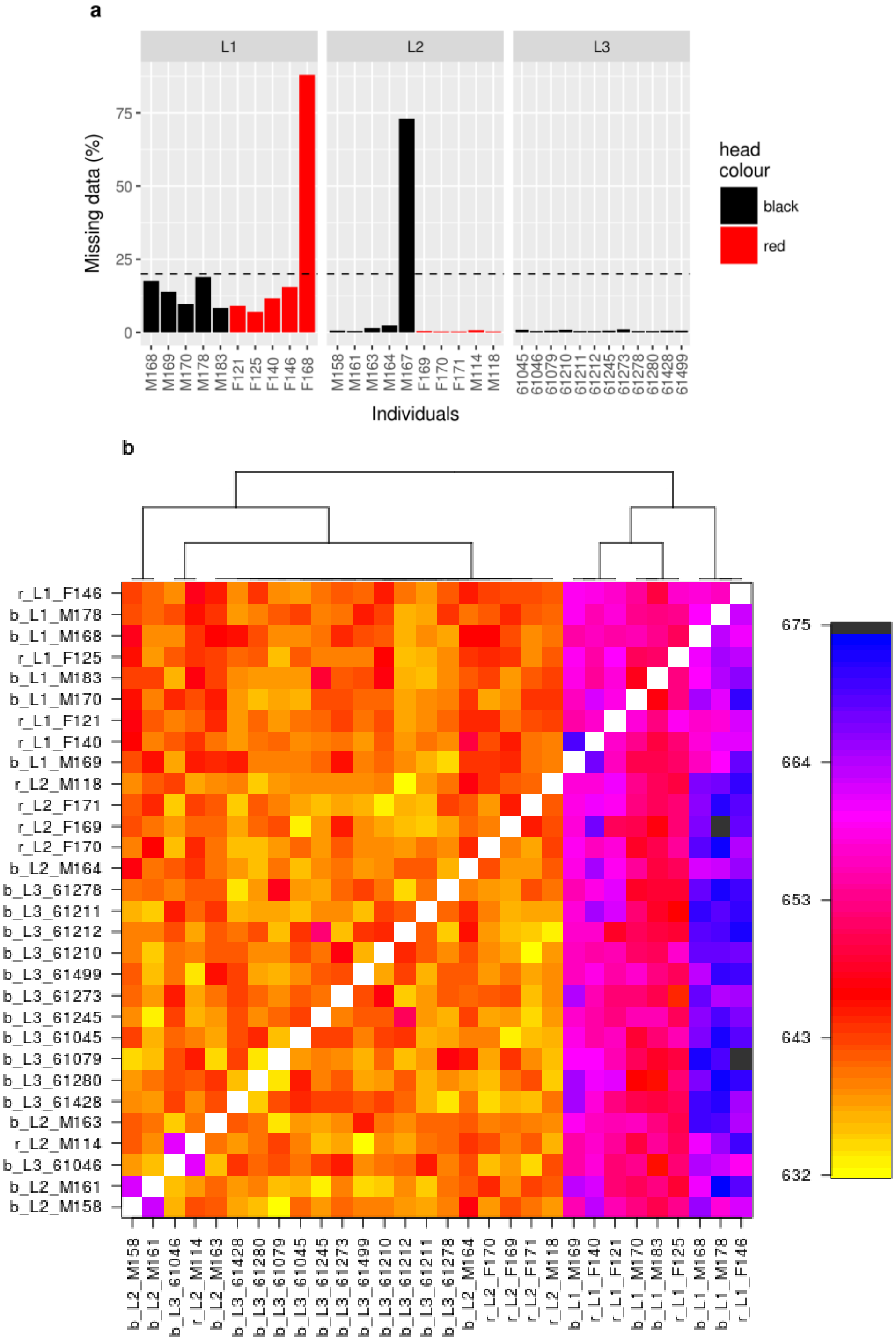


**Supplementary Figure 6 |** Test for population substructure between morphs within a wild population ($n_{black}$ = 22, and $n_{red}$ = 10) using 18,719 autosomal RADSeq loci. **a**, Per-sample missingness in the RADSeq dataset. Samples with > 20% of missing data (dashed line) were excluded from fineRADstructure analysis. **b**, Clustered fineRADstructure coancestry matrix. Head colour (b: black, r: red) and library identity (L1–L3) are added before sample names. Although the variation in the extent of missing data per individual distinguished samples in L1 from the others, there is no apparent cluster that distinguish the morphs.

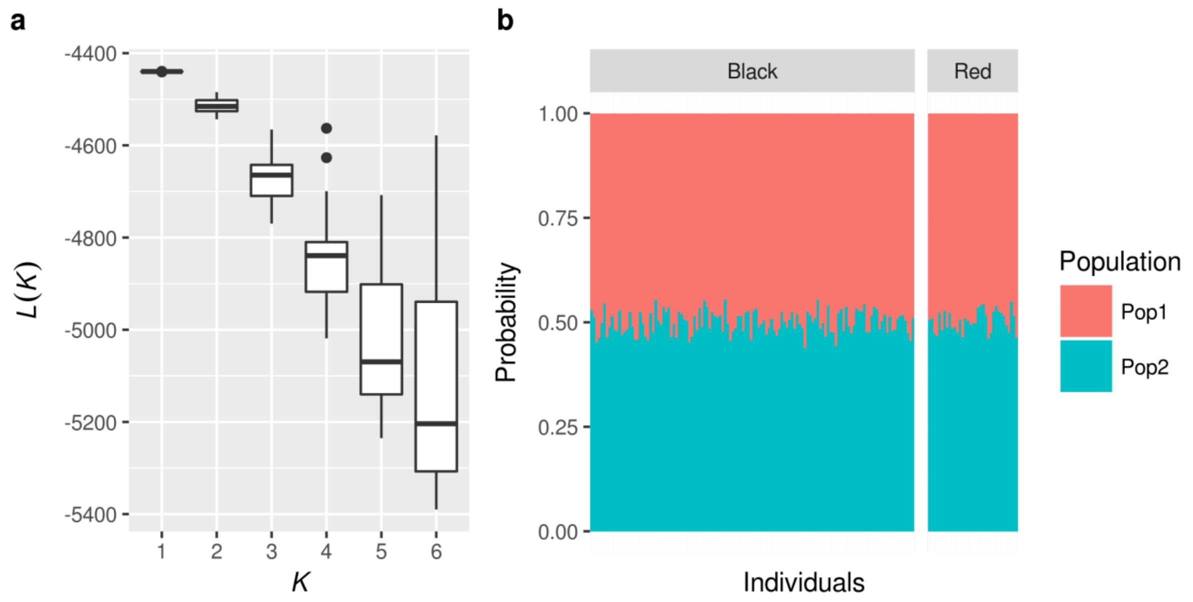


**Supplementary Figure 7 |** Test for population substructure between morphs within a wild population ($n_{black}$ = 126, and $n_{red}$ = 35) using 9 autosomal microsatellite markers. **a**, The log likelihood for each number of hypothetical subpopulations (*K* = 1 to 6) with 20 iterations. **b**, Assignment of individuals to two hypothetical populations**.** Bars represent individual birds, bar colours represent two hypothetical populations, and the proportion of colours within each bar indicates the probability of an individual being assigned to each population.

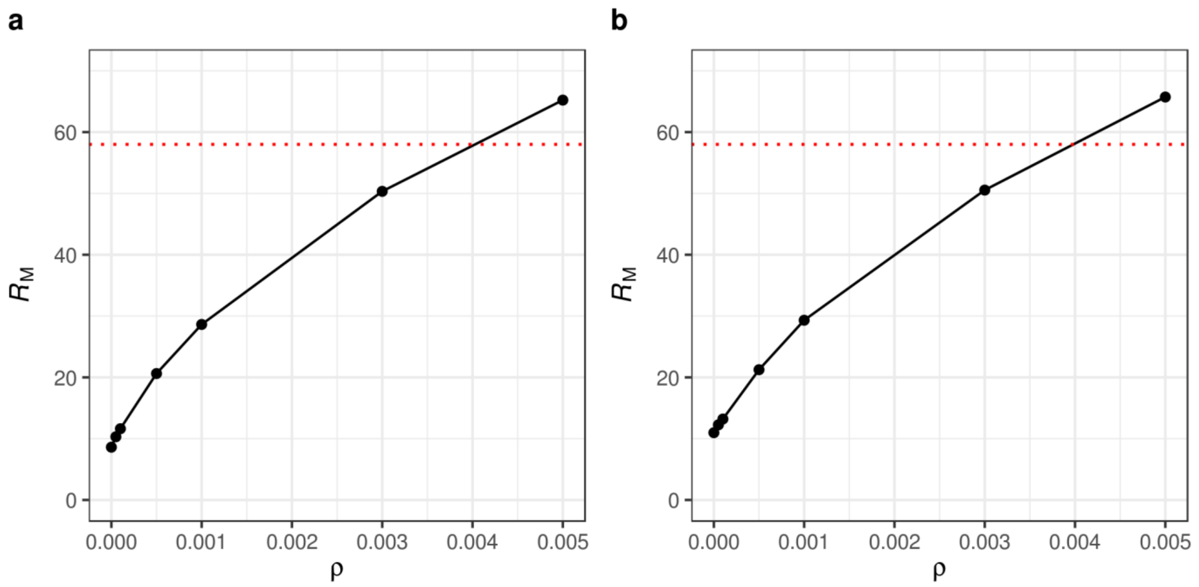


**Supplementary Figure 8 |** Comparison of the observed value of $R_M$ at the 100-kbp sequenced candidate region ($R_M = 58$, red dotted line) with values of $R_M$ obtained from a simulated representation of the sequenced candidate region, for derived allele frequencies of 0.144 (**a**) and 0.856 (**b**), conditioning on a value of $\theta = 1.4416 \times 10^{-2}$, derived from 24 Z-linked intronic reference loci and incorporating a change in population size.

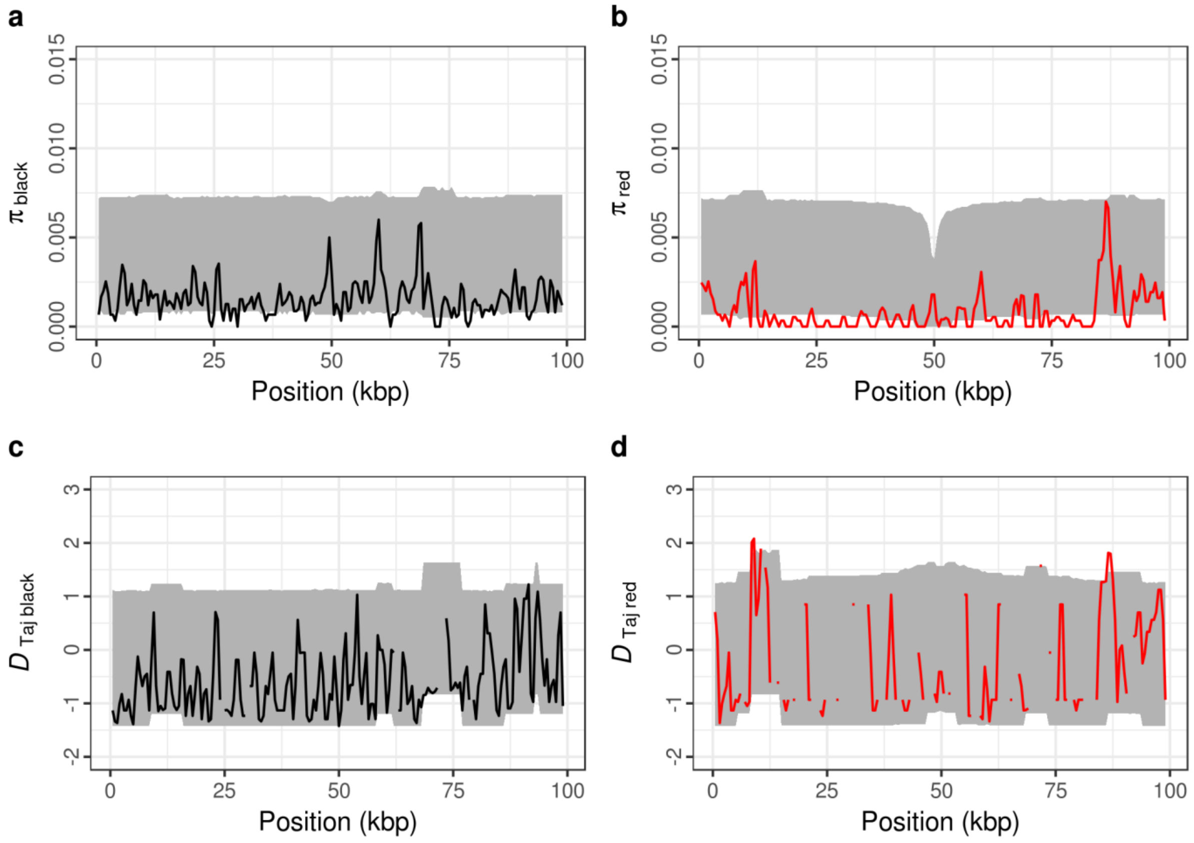


**Supplementary Figure 9 |** A sliding-window comparison of observed within-allelic class polymorphism at the sequenced candidate region with polymorphism simulated under the standard neutral model of molecular evolution incorporating a population size change, with a derived allele frequency of 0.144, taking into account the observed sampling scheme and recombination rate. Solid lines represent the observed data. The grey-shaded regions represent the space encompassed by the 95% confidence intervals derived from the simulations. Position in **a-d** represent ~100-kbp alignment of assembled MiSeq reads for 12 females ($n_{black}$ = 6, $n_{red}$ = 6) as in Fig. 2. **a**, Nucleotide diversity within the black allelic class; **b**, nucleotide diversity within the red allelic class; **c**, Tajima's *D* within the black (ancestral) allelic class; **d**, Tajima's *D* within the red (derived) allelic class. Tajima's *D* was not obtained in a few windows due to a lack of nucleotide diversity.

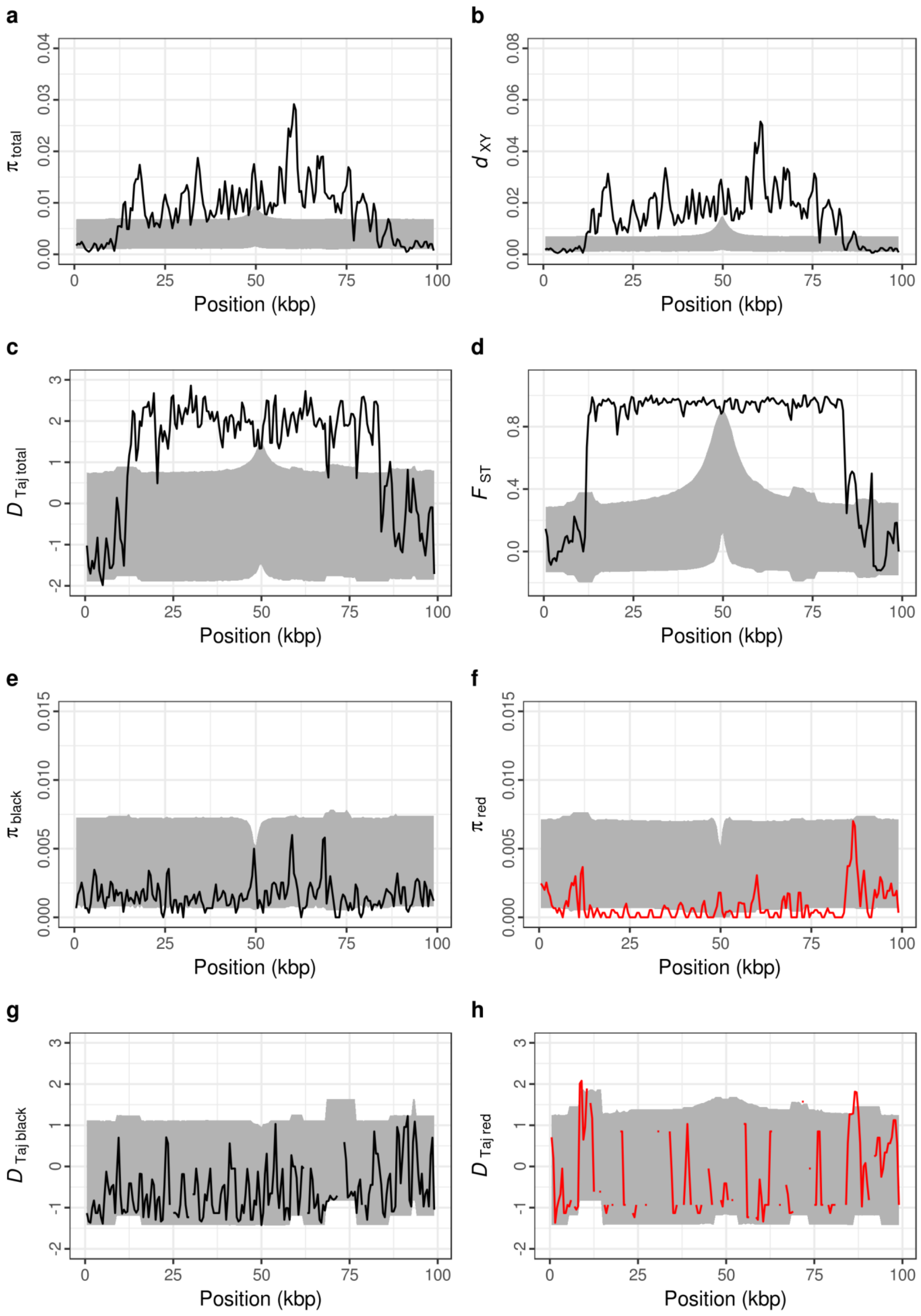


**Supplementary Figure 10 |** A sliding-window comparison of observed between- and within-allelic class polymorphism at the sequenced candidate region with polymorphism simulated under the standard neutral model of molecular evolution incorporating a population size change, with a derived allele frequency of 0.856, taking into account the observed sampling scheme and recombination rate. Solid lines represent the observed data. The grey-shaded regions represent the space encompassed by the 95% confidence intervals derived from the simulations. Position in **a**-**h** represent ~100-kbp alignment of assembled MiSeq reads for 12 females ($n_{black}$ = 6, $n_{red}$ = 6) as in Fig. 2. **a**, Nucleotide diversity for both allelic classes combined; **b**, $d_{XY}$ between both allelic classes; **c**, Tajima's *D* for both allelic classes combined; **d**, $F_{ST}$ between both allelic classes across the candidate region; **e**, nucleotide diversity within the black allelic class; **f**, nucleotide diversity within the red allelic class; **g**, Tajima's *D* within the black (derived) allelic class; **h**, Tajima's *D* within the red (ancestral) allelic class.

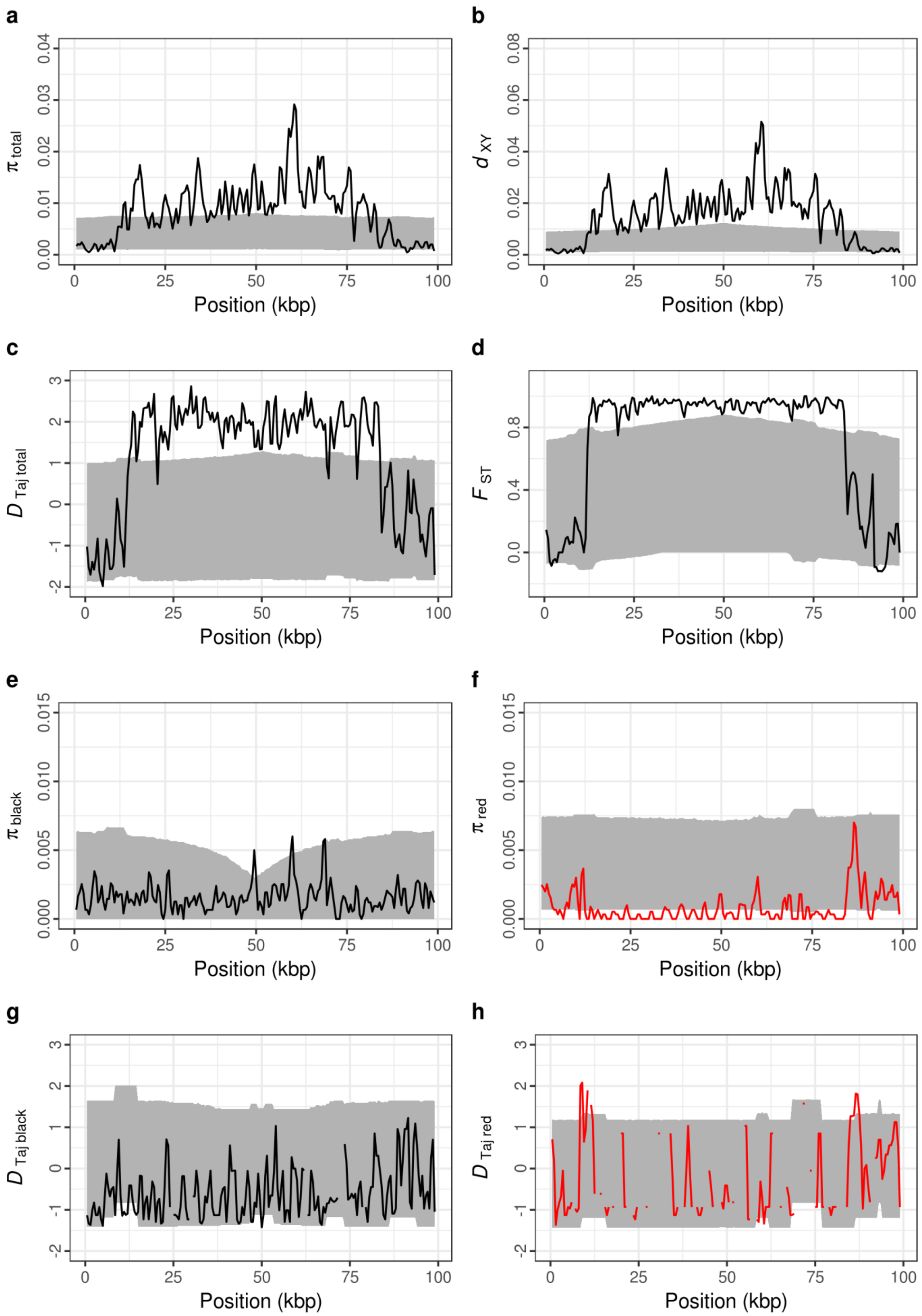


**Supplementary Figure 11 |** A sliding-window comparison of observed between- and within-allelic class polymorphism at the sequenced candidate region with polymorphism simulated under the standard neutral model of molecular evolution incorporating a population size change, with a derived allele frequency of 0.144, taking into account the observed sampling scheme and a reduced population recombination rate $\rho$ = 0.0002. Solid lines represent the observed data. The grey-shaded regions represent the space encompassed by the 95% confidence intervals derived from the simulations. Position in **a**-**h** represent ~100-kbp alignment of assembled MiSeq reads for 12 females ($n_{black}$ = 6, $n_{red}$ = 6) as in Fig. 2. **a**, Nucleotide diversity for both allelic classes combined; **b**, $d_{XY}$ between both allelic classes; **c**, Tajima's *D* for both allelic classes combined; **d**, $F_{ST}$ between both allelic classes across the candidate region; **e**, nucleotide diversity within the black allelic class; **f**, nucleotide diversity within the red allelic class; **g**, Tajima's *D* within the black (ancestral) allelic class; **h**, Tajima's *D* within the red (derived) allelic class.

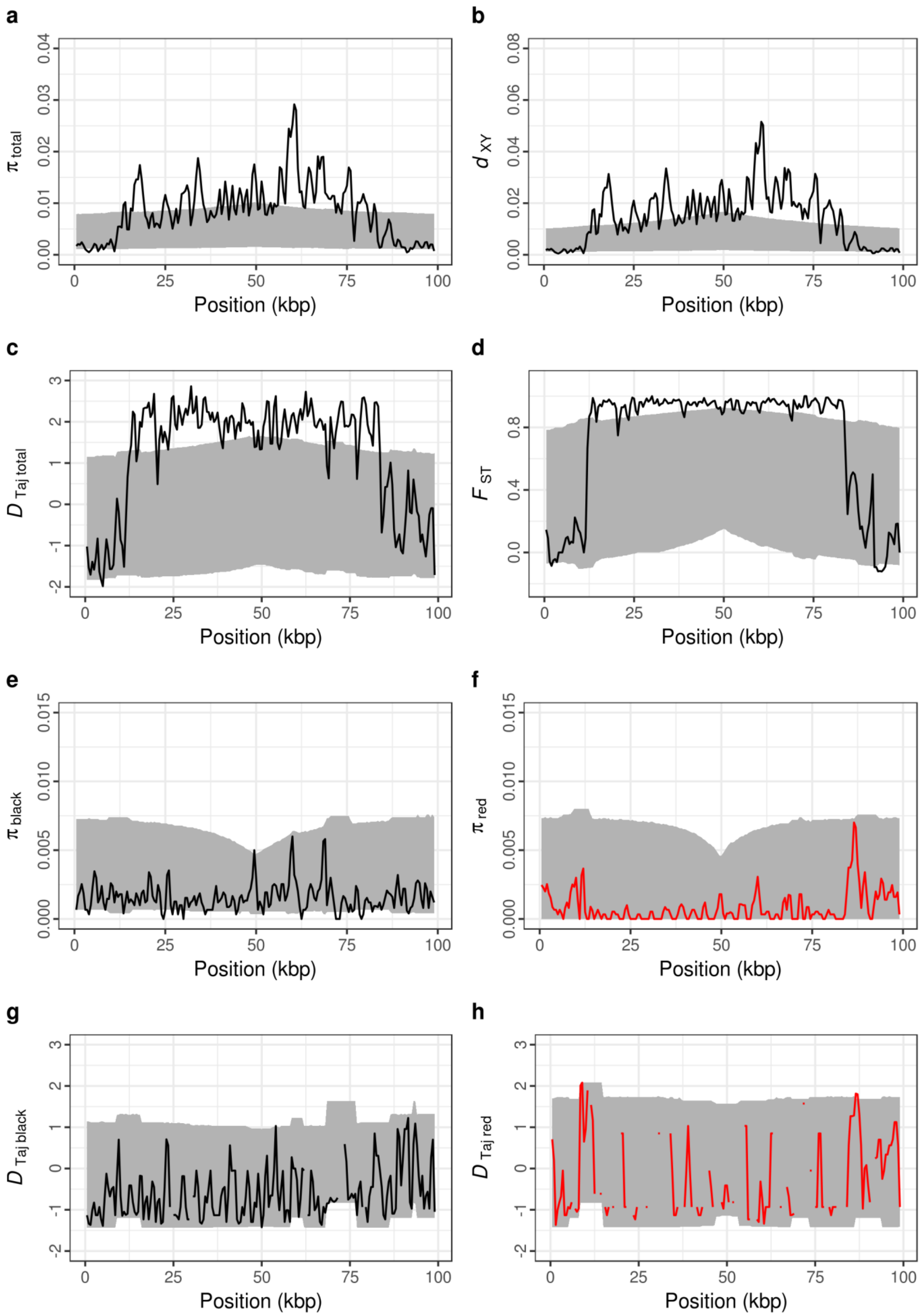


**Supplementary Figure 12 |** A sliding-window comparison of observed between- and within-allelic class polymorphism at the sequenced candidate region with polymorphism simulated under the standard neutral model of molecular evolution incorporating a population size change, with a derived allele frequency of 0.856, taking into account the observed sampling scheme and a reduced population recombination rate $\rho$ = 0.0002. Solid lines represent the observed data. The grey-shaded regions represent the space encompassed by the 95% confidence intervals derived from the simulations. Position in **a**-**h** represent ~100-kbp alignment of assembled MiSeq reads for 12 females ($n_{black}$ = 6, $n_{red}$ = 6) as in Fig. 2. **a**, Nucleotide diversity for both allelic classes combined; **b**, $d_{XY}$ between both allelic classes; **c**, Tajima's *D* for both allelic classes combined; **d**, $F_{ST}$ between both allelic classes across the candidate region; **e**, nucleotide diversity within the black allelic class; **f**, nucleotide diversity within the red allelic class; **g**, Tajima's *D* within the black (derived) allelic class; **h**, Tajima's *D* within the red (ancestral) allelic class.

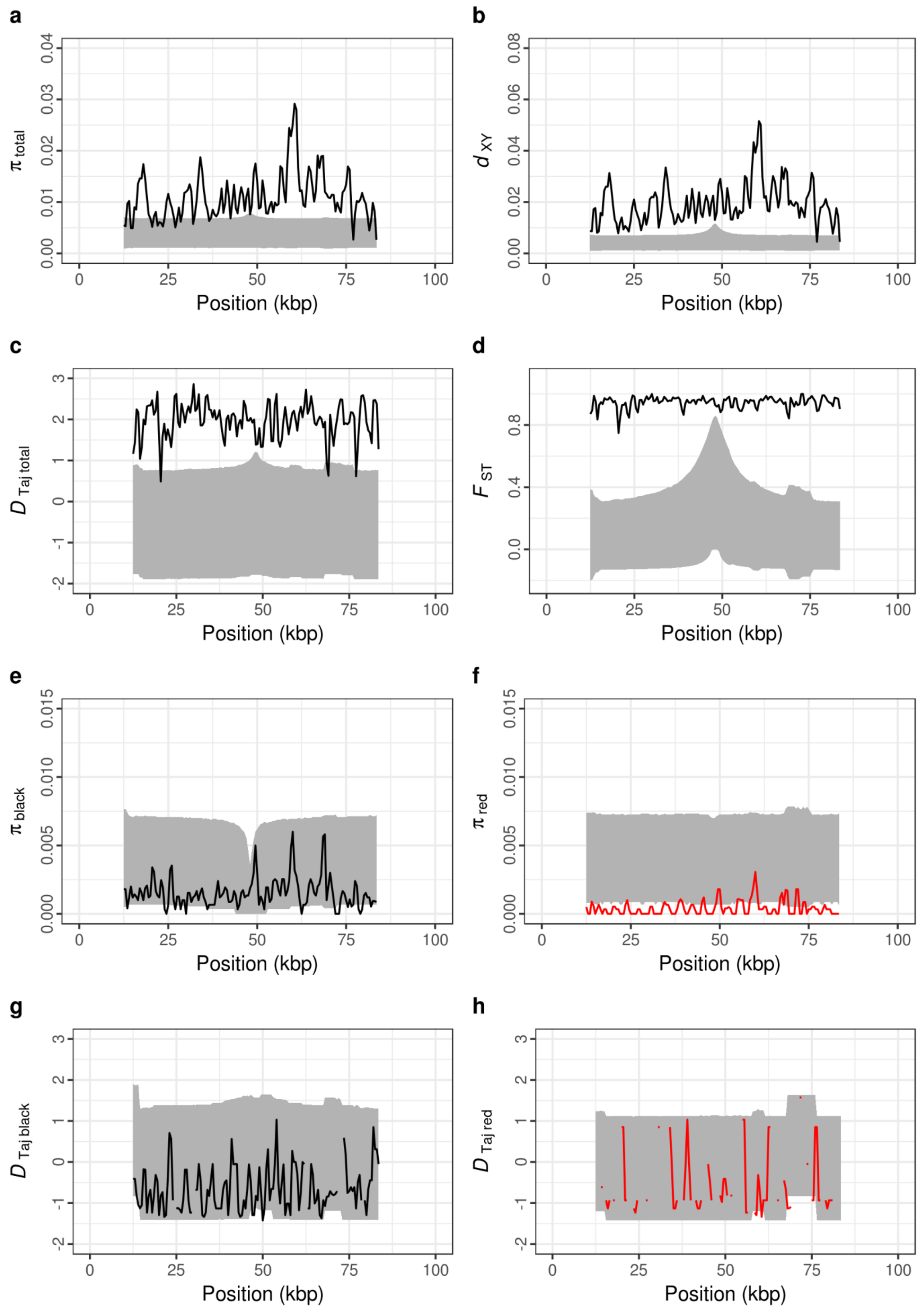


**Supplementary Figure 13 |** A sliding-window comparison of observed between- and within-allelic class polymorphism of the *Red* locus alone (at positions 12–84 kbp of the 100-kbp sequenced candidate region) with polymorphism simulated under the standard neutral model of molecular evolution incorporating a population size change, with a derived allele frequency of 0.144, taking into account the observed sampling scheme and population recombination rate calculated for the reduced-length region. Solid lines represent the observed data. The grey-shaded region represents the space encompassed by the 95% confidence intervals derived from the simulations. Position in **a**-**h** represent positions relative to the full ~100-kbp alignment of assembled MiSeq reads for 12 females ($n_{\mathrm{black}}$ = 6, $n_{\mathrm{red}}$ = 6), as in Fig. 2. **a**, Nucleotide diversity for both allelic classes combined; **b**, $d_{XY}$ between both allelic classes; **c**, Tajima's *D* for both allelic classes combined; **d**, $F_{ST}$ between both allelic classes across the candidate region; **e**, nucleotide diversity within the black allelic class; **f**, nucleotide diversity within the red allelic class; **g**, Tajima's *D* within the black (ancestral) allelic class; **h**, Tajima's *D* within the red (derived) allelic class.

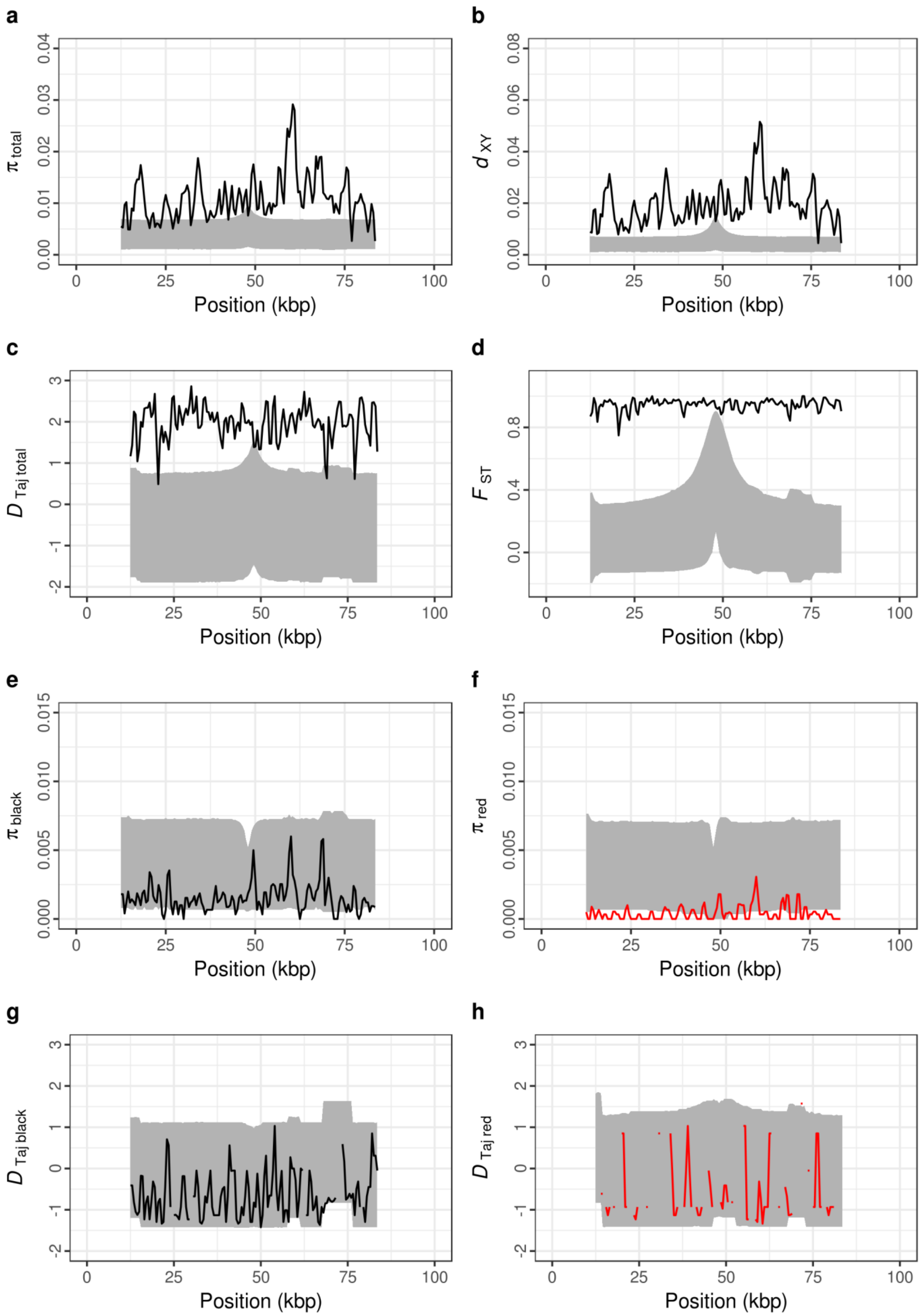


**Supplementary Figure 14 |** A sliding-window comparison of observed between- and within-allelic class polymorphism of the *Red* locus alone (at positions 12–84 kbp of the 100-kbp sequenced candidate region) with polymorphism simulated under the standard neutral model of molecular evolution incorporating a population size change, with a derived allele frequency of 0.856, taking into account the observed sampling scheme and population recombination rate calculated for the reduced-length region. Solid lines represent the observed data. The grey-shaded region represents the space encompassed by the 95% confidence intervals derived from the simulations. Position in **a**-**h** represent positions relative to the full ~100-kbp alignment of assembled MiSeq reads for 12 females ($n_{\text{black}}$ = 6, $n_{\text{red}}$ = 6), as in Fig. 2. **a**, Nucleotide diversity for both allelic classes combined; **b**, $d_{XY}$ between both allelic classes; **c**, Tajima's *D* for both allelic classes combined; **d**, $F_{ST}$ between both allelic classes across the candidate region; **e**, nucleotide diversity within the black allelic class; **f**, nucleotide diversity within the red allelic class; **g**, Tajima's *D* within the black (derived) allelic class; **h**, Tajima's *D* within the red (ancestral) allelic class.

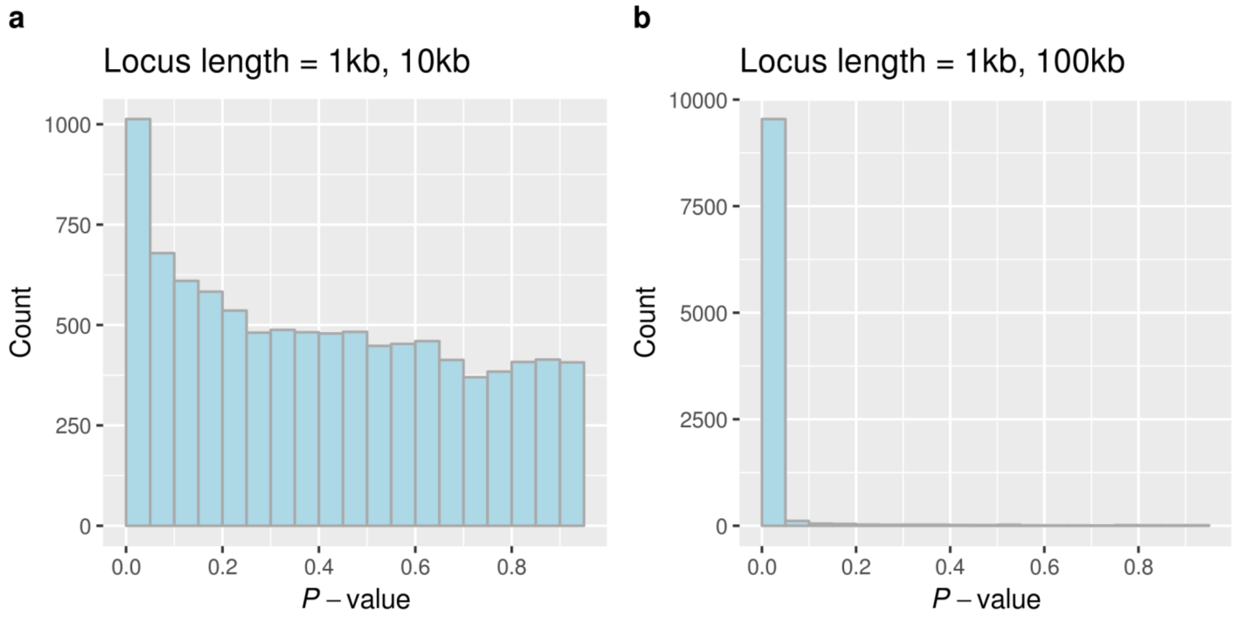


**Supplementary Figure 15 |** The effect of unequal locus lengths on the HKA test's *P*-value distribution, obtained from a $\chi^2$ distribution with 1 d.f.. We simulated two loci with 12 chromosomes under the standard neutral model, with values of $\theta$ for polymorphism and divergence informed by the Gouldian finch–zebra finch system, either **a**, 1 kbp and 10 kbp in length, or **b**, 1 kbp and 100 kbp in length. If data are obtained from equal-sized loci then the *P*-values should be uniformly distributed, as shown in Supplementary Figure 16a. The (greater) surfeit of low *P*-values with (more) unequal locus lengths suggests that the test is anti-conservative when locus lengths are unequal.

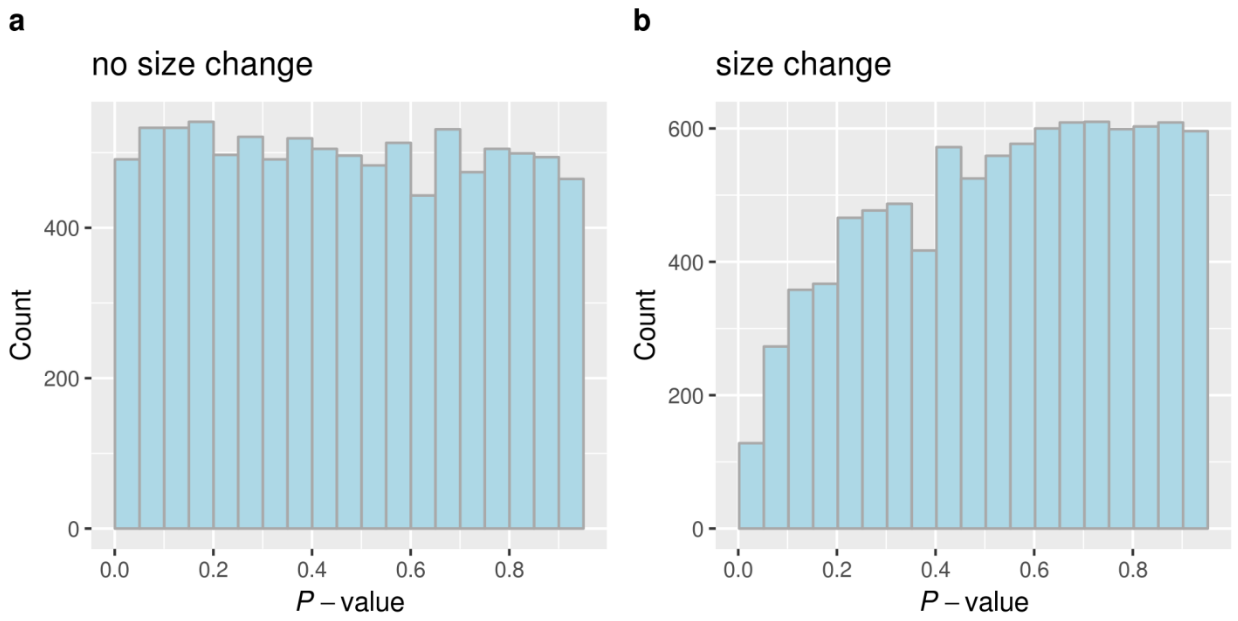


**Supplementary Figure 16 |** The effect of a population size change on the HKA test's *P*-value distribution, obtained from a $\chi^2$ distribution with 1 d.f. **a**, Under no size change, we simulated two 1-kbp loci with 12 chromosomes under the standard neutral model, with values of $\theta$ for polymorphism and divergence informed by the Gouldian finch–zebra finch system. **b**, The size-change model is identical, with the exception that we included the inferred population expansion in the Gouldian finch lineage in our simulations. If data are obtained from the model assumed in the original formulation of the HKA test, the *P*-values should be uniformly distributed, as shown in **a**. The deficit of low *P*-values under a size change model (**b**) suggests that the test should be conservative.

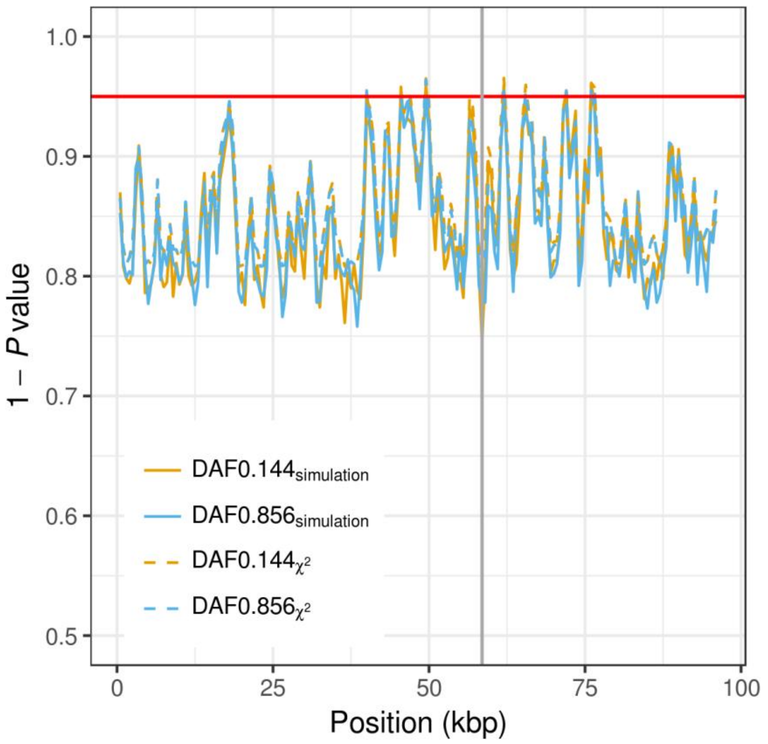


**Supplementary Figure 17 |** Sliding-window implementation of the HKA test, taking into account non-random sampling and recombination. Position represents ~100 kbp alignment of assembled MiSeq reads for 12 females ($n_{black}$ = 6, $n_{red}$ = 6) as in Fig. 2. The region of the *Red* locus incorporating the putative transposable element in the Gouldian finch lineage has been removed, and the position of the focal site set to its position (the vertical grey line). The horizontal red line indicates a *P*-value of 0.05.

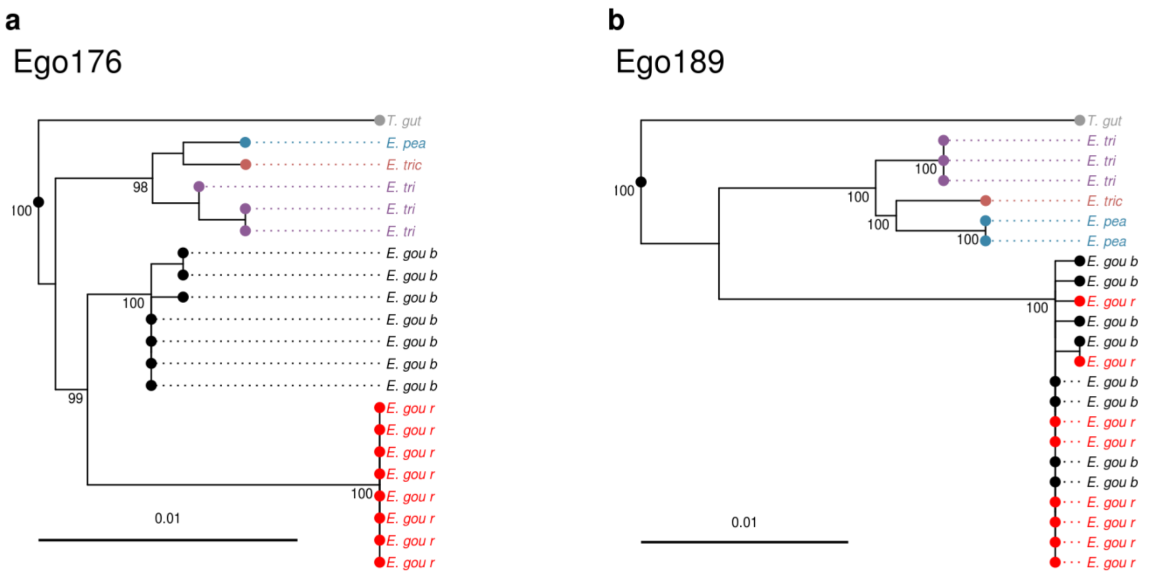


**Supplementary Figure 18 |** Gene genealogy for the *Red* locus and in the flanking region. Maximum-likelihood trees (GTR+Γ+I) for the species in the family *Estrildidae* using sequences for loci **a**, *Ego176* in the *Red* locus and **b**, *Ego189* in the flanking region (See Supplementary Table 4 for the position and the primer sequences for each locus). Abbreviated species names are shown at the tips of branches (*T. gut*: *Taeniopygia guttata*; *E. pea*: *Erythrura pealii*; *E. tri*: *Erythrura tricolor*; *E. tric*: *Erythrura trichroa*; *E. gou b*: *Erythrura gouldiae* black; *E. gou r*: *Erythrura gouldiae* red). Node support values (%) were generated from 1,000 bootstrap replicates and values >80% are shown next to the branches.

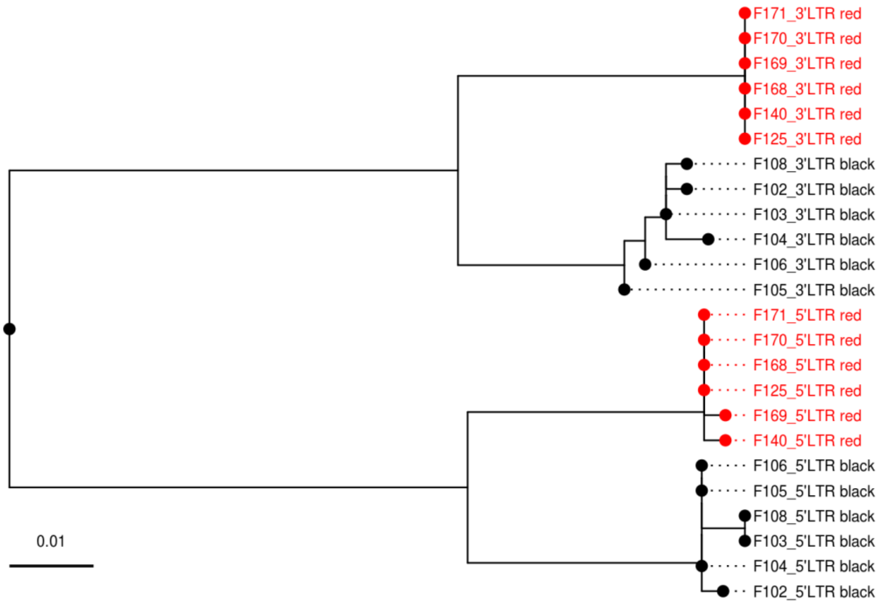


**Supplementary Figure 19 |** Gene genealogy of repeat sequences in LTRs near *M1*. An unrooted maximum-likelihood tree (GTR+Γ+I) of the 3' and 5' repeat sequences at the ends of the putative long terminal repeat (LTR) retrotransposon in the 2.8-kbp insertion identified near the *M1* region that is not present in the zebra finch genome. The inserted region shows the features of an LTR retrotransposon (i.e. an ~400-bp direct repeat sequence at each end and identical (CTACAT) flanking sequences at each end of the insertion) and is found in several places in the zebra finch genome. LTRs have identical repeat sequences at the time of insertion, which diverge subsequently. If insertion of the retrotransposon initiated the colour polymorphism, then the sequence divergence between the 3' and 5' repeats should be the same as that between the colour-associated haplotypes. However, much deeper internal branches separated the LTR repeats than those between morphs, suggesting that the polymorphic haplotypes arose long after the insertion event.

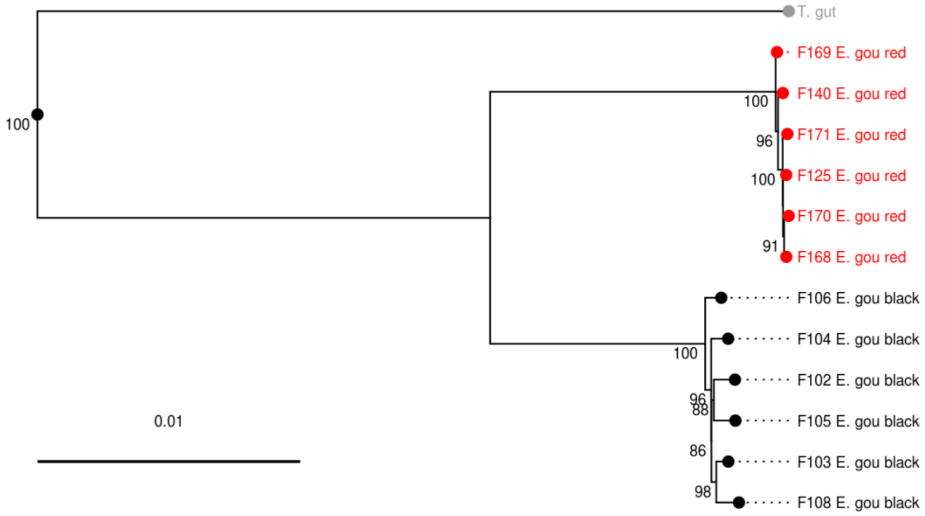


**Supplementary Figure 20 |** Gene genealogy of the *Red* locus. A maximum-likelihood tree using the GTR+Γ+I substitution model was constructed for the *Red* locus (~72 kbp) that showed the highest $F_{ST}$ between the black and red haplotypes. Zebra finch was used as an outgroup. All pairs of black and red sequences showed significant differences in Tajima's nonparametric relative rate test ($P$ < 0.05). Node support values (%) were generated from 1,000 bootstrap replicates and values >80% are shown next to the branches.

**Supplementary Table 1 |** Inter-morph differences between black and red Gouldian finches associated with the *Red* locus, mostly based on observations in captive populations.

| Trait | Black | Red | References |
|---|---|---|---|
| pigmentation | melanin | carotenoids | 11, 12 |
| feather structure | barbs with barbules | no barbules | 12 |
| social dominance | submissive | dominant | 13, 14 |
| personality | bolder, risk taker | aggressive | 15, 16 |
| stress response | low | high | 17, 18 |
| plasticity in sperm morphometry | low | high | 19 |
| mate choice | assortative | assortative | 19, 20, 21 |
| genetic incompatibility | high | high | 22 |
| sex allocation & maternal investment | overproduction of sons when paired with males of alternative colour morphs | | 19, 23 |

**Supplementary Table 2** | Primers for genotyping the *Red* locus. Primers for *M1* were designed to produce both a 3-bp size and a fluorescence colour difference between alleles in a multiplex PCR. Haplotype-specific primer sequences are underlined. See Methods for the PCR conditions. * Positions are based on zebra finch reference genome.

| Loci | Type of variation* | Primer name | Sequence 5' → 3' |
|---|---|---|---|
| *M1* | 2 SNPs | Ego172_Fa | GAGTTGGAAGCAACCCATGA |
| | 46,561,597–46,561,598 bp | Ego172_Rred | [HEX]TCAGGAGAGCTACATTTAACCTCTTTG |
| | | Ego172_Rblack | [FAM]GGAGAGCTACATTTAACCTCTTCA |
| *M2* | 12-bp indel | Ego173FSd | CAACAGTCACTTCTCATTGTTGTC |
| | 46,522,516–46,522,527 bp | Ego173R_HEXa | [5HEX]AAGAAGCTTGGCCAAAGGAC |
| *M3* | 1 SNP | Ego181Fa | TGTGATTTGAGGTGTTCCAAA |
| | 46,583,055 bp | Ego181R_HEXG | [5HEX]TGCTGATCTGAATGGGTAAGC |
| | | Ego181R_HEXT | [5HEX]AAAGTGCTGATCTGAATGGGTAAGA |

**Supplementary Table 3** | Genotype and gene frequencies at *M1* in the *Red* locus in the Wyndham population. $Z^R$ and $Z^r$ represent the red- and black-linked alleles, respectively. We used Graffelman–Weir's exact test for sex-linked markers to test for the deviation of genotypes of all samples from HWE and obtained the probability using a permutation test (see Methods). Allele frequencies in males and females were not significantly different (Fisher's exact test, $P$ = 0.69). See Supplementary Methods for the maximum-likelihood estimates of allele frequencies.

| | **Number of individuals** | | | | | | |
|---|---|---|---|---|---|---|---|
| | Male | | | | Female | | |
| | $Z^RZ^R$ | $Z^RZ^r$ | $Z^rZ^r$ | Total | $Z^R$ | $Z^r$ | Total |
| Observed | 0 | 27 | 62 | 89 | 9 | 63 | 72 |
| Expected | 2.05 | 22.90 | 64.05 | 89 | | | |
| | Graffelman-Weir's exact test, $P_{\text{permutation}}$ = 0.24 | | | | | | |
| | **Number of genes** | | | | **Allele frequencies** | | |
| | $Z^R$ | $Z^r$ | Total | | $Z^R$ | $Z^r$ | |
| Males | 27 | 151 | 178 | | 0.85 | 0.15 | |
| Females | 9 | 63 | 72 | | 0.88 | 0.12 | |

**Supplementary Table 4 |** Primers used in the test for the extent of LD around the *Red* locus. Amplicons were sequenced using the Sanger method. Gene names and the (I)ntron and (E)xon positions sequenced are indicated under Target. The positions are based on the zebra finch reference genomic sequence. Approximate position of the *Red* locus is shaded.

| Loci | Target | PCR start | PCR End | Forward primers (5'→ 3') | Reverse primers (5'→ 3') |
|---|---|---|---|---|---|
| **Control loci** | | | | | |
| Ego31 | SMAD2; I6 | 414,579 | 415,743 | TCTCCAGCAGAGCTGTCTCC | ACGCAGGCTCCGAGTAAGTA |
| Ego30 | SMAD2; I2 | 421,173 | 422,371 | CAGCAGGCCTTTACAGCTTC | CGGTGAGACACCTGCAGAC |
| Ego35 | SLC45A2; I5 | 41,046,360 | 41,047,343 | AGGGGTAGAGATGGGATGCT | TCCAGTACCCAGTCCAAAAAG |
| Ego34 | SLC45A2; I1 | 41,056,258 | 41,057,392 | TACCTCAATGGGGACGTGAT | GTCCGCTGCTTGTCTCTCTC |
| Ego66 | SFRS12; I12 | 51,193,251 | 51,194,448 | TGACAAAAGCACCACTACTTTGA | TTCCCATTCTGTTGTAGCTTGA |
| Ego67 | PIK3R1; I10 | 51,950,868 | 51,951,996 | GCATTGAAGCAGTAGGAAAGAAA | AAATGCCTCAATGGCTGTTC |
| Ego68 | GOLM1; I3 | 52,030,810 | 52,031,841 | GATCGATTGATTCAGAGTTTGC | CATCCAGCTGTAGCTTTCCA |
| Ego10 | SLC28A3; I8E9 | 52,630,547 | 52,631,503 | CTGGCCTGAAACTGAAAAGC | CAAGAAAGGGCTGAGTCGAG |
| Ego28 | intergenic | 53,371,312 | 53,372,310 | GGCCCTAAAATAGCATGAGAA | CAGCATTTCAGCAGAAGCAA |
| Ego9 | GNAQ; I4 | 54,782,996 | 54,783,940 | CAAAGCGTCATTTTCAGGTAGG | ATCAAGTCCATGAACACCTTGA |
| Ego55 | PAR2; I1 | 56,982,536 | 56,983,699 | AGCCTGTTAGCAGAGGAGGA | GCCAATGCCAAGTTAACCAT |
| Ego43 | TYRP1; I6 | 60,967,893 | 60,968,642 | GAGAATGCCCCTATTGGACA | ATGCAGCAGCAGCAAAGATA |
| Ego42 | TYRP1; I3 | 60,971,686 | 60,972,668 | TCTCTCAGTGGCGAGTGCTA | TCGTGCTACATTTCCAGCAG |
| Ego44 | FSD1L; I2 | 67,712,034 | 67,713,008 | CGCAACTTCATTGACATGCT | AGCTCACTGATGGCATGAAA |
| Ego45 | FSD1L; I10 E11 I11 | 67,727,068 | 67,728,167 | TTGCTGGTGAATCCTACACA | AATGCCTTTGAGCTTGCATT |
| ***Red*** **locus** | | | | | |
| Ego199 | MOCS2; I4 | 46,458,425 | 46,459,349 | GCCAGTGCACCAGTAAATGA | TGGCGCATCTTCTCTTTCTT |
| Ego189 | Intergenic | 46,491,656 | 46,492,712 | GCAGCATGTAAATCAGCATCA | CAGAACAATGCTGGGATCAA |
| Ego190 | Intergenic | 46,497,744 | 46,498,649 | CAGTGAGGGGACAGCAAATC | GCACTGCAGGAGTGTGGTAA |
| Ego192 | Intergenic | 46,508,699 | 46,509,718 | TGGTTCACAAGCTGCATTTC | GGCACGTAAAAGGGTTTTTG |
| Ego194 | Red | 46,517,972 | 46,519,034 | CGCCCAATGAGTTATTGACA | CTCTGTGCCAGCCATATGAA |
| Ego173 | Red | 46,522,336 | 46,523,362 | CAACTTAGGACTGGAACAACTCA | CCCTCCACAAGAATGCAAAT |
| Ego195 | Red | 46,527,693 | 46,528,731 | CAGGCTGGACAAATCAGGTT | TGCCATGTGGGGTTTTTATT |
| Ego196 | Red | 46,532,186 | 46,533,192 | GCCTGCTATTGATTTTACATTCC | AATGCCTGAAAGCAGTTTGAA |
| Ego197 | Red | 46,536,764 | 46,537,675 | GGTGGGTGTCAGGCTGTATT | GCCATGCTCTGTCCCTAAAC |
| Ego198 | Red | 46,541,858 | 46,542,787 | TCGTGTACCTCCACCATTCA | TGCTGCTAGTTCCACGAGAG |
| Ego175 | Red | 46,546,968 | 46,547,927 | AGTCCAGCCATTCTGCTTGT | CCACACATTGGTTTTCATGC |
| Ego176 | Red | 46,550,604 | 46,551,556 | AACACTCAAGTATTGTGCAAGAAAA | CCCTGGAGGTTCCACAAGTA |
| Ego165 | Red | 46,561,114 | 46,562,005 | TCCTTCCTGCTGTCCTTGAC | TGGATGAGGCCTTCTACAGG |
| Ego166 | Red | 46,565,160 | 46,566,036 | ACCACCTTGTTCCCCAAATC | AGCTTCATTCCTCTGGGGTTA |
| Ego178 | Red | 46,570,267 | 46,571,078 | CGAGGGCAAGTCAGGTTATT | CCCTGATTTGTGGAGACACC |
| Ego180 | Red | 46,577,986 | 46,578,792 | AATTCCGACCCACTTGTCTG | TCCTTTGGTGTTTTTGTAACAATG |
| Ego181 | Red | 46,582,953 | 46,583,736 | TGTGATTTGAGGTGTTCCAAA | GCCACAAAAGAAATCTGTCCA |
| Ego182 | Intergenic | 46,588,126 | 46,589,120 | GCACAGGAACAAACTGCAAA | AAGCCAAATGCATTCAGTCC |
| Ego184 | Intergenic | 46,600,895 | 46,601,744 | CAGGAATTTGCTTGTTTTTCAC | GGCTGGCCTCTGAATAAAAT |
| Ego167 | FST 5' UTR & E1 | 46,604,796 | 46,605,396 | CAGAGCCTGACGTTCATTCA | CAGCTGTAATCCTGGGGTTC |
| Ego168 | FST E2 | 46,606,840 | 46,607,536 | TTCCAAATGGGCCGTAGTAG | TGTGCAGAAGAGACCTGGTG |
| Ego38 | FST E3 | 46,607,166 | 46,608,164 | AAACTGCATCCCATGCAAAG | CTGCCTGGGCATAAGACATC |
| Ego169 | FST E4 | 46,608,003 | 46,608,625 | GTCATCCAGCAGCCCTTAAA | GGGAGGTTGTTTCTCCATTG |
| Ego170 | FST E5 | 46,609,228 | 46,609,854 | GCACTGTTGTCCAAGGGAAT | CCCCAAAAACTCCAACTTGA |
| Ego39 | FST I5 | 46,609,556 | 46,610,404 | ACTTACCCAAGCGAGTGTGC | CCTGGTCTTCATCTTCCTCTTC |
| Ego171 | FST E6 & 3' UTR | 46,610,202 | 46,610,918 | GCATAACAGCAGATGCCAAA | TGCCTTTTGGTCAGTCAAGA |

**Supplementary Table 5 |** Fixed and shared polymorphisms in 6–8 each of black and red females at the control loci and the *Red* locus. See Supplementary Table 4 for the position of loci. Approximate position of the *Red* locus is shaded.

| Locus | $L$ (bp) | $n_{black}$ | $n_{red}$ | $S_{black}$ | $S_{red}$ | $S_{fixed}$ | $S_{shared}$ |
|---|---|---|---|---|---|---|---|
| **Control loci** | | | | | | | |
| Ego31 | 941 | 8 | 8 | 1 | 2 | 0 | 4 |
| Ego30 | 1005 | 8 | 8 | 3 | 5 | 0 | 2 |
| Ego35 | 830 | 8 | 8 | 2 | 3 | 0 | 7 |
| Ego34 | 978 | 8 | 8 | 1 | 3 | 0 | 3 |
| Ego66 | 1022 | 8 | 7 | 2 | 3 | 0 | 0 |
| Ego67 | 913 | 8 | 8 | 5 | 10 | 0 | 5 |
| Ego68 | 900 | 8 | 7 | 2 | 1 | 0 | 8 |
| Ego10 | 758 | 8 | 8 | 6 | 0 | 0 | 8 |
| Ego28 | 796 | 8 | 8 | 4 | 4 | 0 | 4 |
| Ego9 | 733 | 8 | 8 | 4 | 4 | 0 | 0 |
| Ego55 | 815 | 8 | 8 | 1 | 3 | 0 | 15 |
| Ego43 | 529 | 8 | 8 | 1 | 6 | 0 | 9 |
| Ego42 | 795 | 8 | 8 | 2 | 10 | 0 | 12 |
| Ego44 | 786 | 8 | 7 | 1 | 5 | 0 | 6 |
| Ego45 | 568 | 8 | 8 | 1 | 3 | 0 | 3 |
| ***Red* locus** | | | | | | | |
| Ego199 | 681 | 8 | 7 | 4 | 2 | 0 | 0 |
| Ego189 | 788 | 8 | 8 | 3 | 1 | 0 | 1 |
| Ego190 | 729 | 8 | 8 | 1 | 3 | 0 | 6 |
| Ego192 | 829 | 8 | 8 | 8 | 1 | 0 | 0 |
| Ego194 | 881 | 8 | 8 | 4 | 0 | 6 | 0 |
| Ego173 | 812 | 8 | 8 | 5 | 0 | 9 | 0 |
| Ego195 | 858 | 8 | 8 | 3 | 1 | 14 | 0 |
| Ego196 | 791 | 8 | 8 | 1 | 0 | 9 | 0 |
| Ego197 | 706 | 8 | 7 | 5 | 0 | 8 | 0 |
| Ego198 | 800 | 8 | 8 | 4 | 1 | 8 | 0 |
| Ego175 | 771 | 8 | 8 | 3 | 1 | 12 | 0 |
| Ego176 | 815 | 7 | 8 | 2 | 0 | 11 | 0 |
| Ego165 | 720 | 8 | 8 | 6 | 0 | 20 | 0 |
| Ego166 | 713 | 8 | 8 | 4 | 0 | 14 | 0 |
| Ego178 | 602 | 8 | 8 | 10 | 1 | 16 | 0 |
| Ego180 | 725 | 8 | 8 | 2 | 1 | 14 | 0 |
| Ego181 | 695 | 8 | 8 | 2 | 1 | 5 | 0 |
| Ego182 | 871 | 8 | 8 | 2 | 11 | 0 | 1 |
| Ego184 | 709 | 8 | 7 | 4 | 2 | 0 | 0 |
| Ego167 | 562 | 8 | 8 | 1 | 0 | 0 | 1 |
| Ego168 | 613 | 8 | 8 | 5 | 1 | 0 | 3 |
| Ego38 | 861 | 8 | 8 | 3 | 3 | 0 | 4 |
| Ego169 | 599 | 8 | 7 | 0 | 0 | 0 | 2 |
| Ego170 | 591 | 8 | 8 | 4 | 0 | 0 | 1 |
| Ego39 | 699 | 7 | 8 | 0 | 1 | 0 | 0 |
| Ego171 | 573 | 7 | 6 | 0 | 0 | 0 | 0 |

$L$, total number of sites excluding gaps. $n_{black}$ and $n_{red}$ refer to the numbers of chromosomes sequenced from black and red females, respectively. $S$-values indicate numbers of variable sites: numbers of SNPs exclusive to black ($S_{black}$) or red ($S_{red}$), fixed differences between chromosomes ($S_{fixed}$) or shared between chromosomes ($S_{shared}$).

**Supplementary Table 6 |** Primer sets for the long-range PCR used in the test for the presence of inversions around the *Red* locus. The amplicons were sequenced on an Illumina MiSeq sequencer and the total ~100-kbp sequence for each allele was used in the simulations. The start and end position of each amplicon is based on the reference sequence of the zebra finch. Approximate position of the *Red* locus is shaded.

| Forward primer | Forward primers (5’→ 3’) | Reverse primer | Reverse primers (5’→ 3’) | PCR Start | PCR End | Overlap* |
|---|---|---|---|---|---|---|
| Ego191_Fb | AGTGGCAGGAACAGTGTGTG | Ego192_Rb | GGCACGTAAAAGGGTTTTTG | 46,503,706 | 46,509,718 | 1,020 |
| Ego192_Fb | TGGTTCACAAGCTGCATTTC | Ego193_Ra | GCAAGAACTAAGGGCCAACA | 46,508,699 | 46,512,564 | 517 |
| Ego193_FSa | TTGGTGGCTTAGTGGTGTATTG | Ego194_RSa | ACTTCCACCAAAACAGATGTCT | 46,512,048 | 46,518,757 | 786 |
| Ego194_Fb | CGCCCAATGAGTTATTGACA | Ego173_Rb | CCCTCCACAAGAATGCAAAT | 46,517,972 | 46,523,362 | 587 |
| Ego173_FSc | AATTGCAGCATTAGGCTTGG | Ego195_RSa | TCTTGTGCTTTCTCTGGCATT | 46,522,776 | 46,528,453 | 761 |
| Ego195_Fb | CAGGCTGGACAAATCAGGTT | Ego196_Rb | AATGCCTGAAAGCAGTTTGAA | 46,527,693 | 46,533,192 | 1,007 |
| Ego196_Fb | GCCTGCTATTGATTTTACATTCC | Ego197_Ra | GCCATGCTCTGTCCCTAAAC | 46,532,186 | 46,537,675 | 912 |
| Ego197_Fb | GGTGGGTGTCAGGCTGTATT | Ego198_Rb | TGCTGCTAGTTCCACGAGAG | 46,536,764 | 46,542,787 | 503 |
| Ego198_FSb | TGCTTTATTCTGGAGTTTGAACC | Ego175_RSa | TCCACTCTTACCTCTGGACTTC | 46,542,285 | 46,547,758 | 791 |
| Ego175_Fa | AGTCCAGCCATTCTGCTTGT | Ego176_Rb | CCCTGGAGGTTCCACAAGTA | 46,546,968 | 46,551,556 | 827 |
| Ego176_FSb | CCACCCAGGTAAATGACTCATG | Ego177_RSc | GTCATTGCAGCTGTGTTGGA | 46,550,730 | 46,558,040 | 455 |
| Ego177_Fb | ATGTTTTCATGGCCTGCTTC | Ego165_LRc | TGGATGAGGCCTTCTACAGG | 46,557,586 | 46,562,005 | 892 |
| Ego165_Fd | TCCTTCCTGCTGTCCTTGAC | Ego166_Rb | AGCTTCATTCCTCTGGGGTTA | 46,561,114 | 46,566,036 | 877 |
| Ego166_Fa | ACCACCTTGTTCCCCAAATC | Ego178_Ra | CCCTGATTTGTGGAGACACC | 46,565,160 | 46,571,078 | 812 |
| Ego178_Fa | CGAGGGCAAGTCAGGTTATT | Ego180_Ra | TCCTTTGGTGTTTTTGTAACAATG | 46,570,267 | 46,578,792 | 755 |
| Ego180_FSa | TGCAAACACCAAATATGCAG | Ego181_RSb | ATGCCAGCAGTGTGTTAAGC | 46,578,038 | 46,583,348 | 202 |
| Ego181_FSa | TACCTGGGCTTGTGACAACA | Ego182_RSb | ACATGTACTTGTGGGATAGTTGA | 46,583,147 | 46,589,020 | 471 |
| Ego182_FSa | TCGTCCTTGCAGTGTGAGTTAGAA | Ego183_RSc | ACAGTCTAACTTGCTGTGTCTGGA | 46,588,550 | 46,595,428 | 136 |
| Ego183_Fa | CTGCTGCAGGATAGAAAGAAG | Ego184_Ra | GGCTGGCCTCTGAATAAAAT | 46,595,293 | 46,601,744 | |

*Length of sequence overlapping with that in the next row.

**Supplementary Table 7 |** Success of long-range PCR across the *Red* locus for hemizygous females ($n_{black}$ = 6, $n_{red}$ = 6), showing that there was no evidence of an inversion between the morphs. All primers amplified except those denoted as NA (Not Amplified). See Supplementary Table 6 for primer sequences and their locations. Approximate position of the *Red* locus is shaded.

| Primers | | Black | | | | | | Red | | | | | |
|---|---|---|---|---|---|---|---|---|---|---|---|---|---|
| Forward | Reverse | F102 | F103 | F104 | F105 | F106 | F108 | F125 | F140 | F168 | F169 | F170 | F171 |
| Ego191_Fb | Ego192_Rb | | | | | | | | | | | | |
| Ego192_Fb | Ego193_Ra | | | | | | | | **NA** | | | | |
| Ego193_FSa | Ego194_Rsa | | | **NA** | | | | | **NA** | | | **NA** | |
| Ego194_Fb | Ego173_Rb | | | **NA** | | | | | | | | | |
| Ego173_FSc | Ego195_RSa | | | | | | | | | | | | |
| Ego195_Fb | Ego196_Rb | | | | | | | | | | | | |
| Ego196_Fb | Ego197_Ra | | | | | | | | | | | | |
| Ego197_Fb | Ego198_Rb | | | | | | | | | | | | |
| Ego198_FSb | Ego175_Rsa | | | | | | | | | | | | |
| Ego175_Fa | Ego176_Rb | | | | | | | | | | | | |
| Ego176_FSd | Ego177_RSc | | | | | | | | **NA** | | | | |
| Ego177_Fb | Ego165_LRc | | | | | | | | | | | | |
| Ego165_Fd | Ego166_Rb | | | **NA** | | | | | | | | | |
| Ego166_Fa | Ego178_Ra | | | | | | | | | | | | |
| Ego178_Fa | Ego180_Ra | | | **NA** | | **NA** | | | | | | **NA** | |
| Ego180_FSa | Ego181_RSb | | | | | | | | | | | | |
| Ego181_FSa | Ego182_RSb | | | | | | | | | | | | |
| Ego182_Fsa | Ego183_RSc | | | **NA** | | | | | **NA** | | | | |
| Ego183_Fa | Ego184_Ra | **NA** | | | | | | | | | | | |

**Supplementary Table 8 |** Twenty-four primer sets for the 'reference loci', designed from the zebra finch Z chromosome reference sequence. The amplicons were sequenced using an Illumina MiSeq sequencer and the assembled sequences were used in the simulations.

| Locus | Target | PCR start | PCR end | Forward primers (5'→ 3') | Reverse primers (5'→ 3') |
|---|---|---|---|---|---|
| Ego210 | KATNAL2 | 87,586 | 88,700 | CTGGGTGTACAGCAGAACCT | TGCTGACTCCTGAGATGCTT |
| Ego213 | LPL | 4,547,064 | 4,548,246 | GGCTGGTTTGTCTTGGTCAT | TGAAAACCCGTGCTCAGATG |
| Ego214 | ADAMTS19 | 5,276,909 | 5,277,903 | CAGATGCTGCCAGAAGATTTATT | AGGACAAGTTTAGTCACCCGA |
| Ego216 | CNTNAP4 | 8,448,096 | 8,449,229 | TGATGGACAGCTACAACATGTG | TGCATTCAATTCTGTGCCTGA |
| Ego217 | DAPK1 | 10,208,734 | 10,209,761 | CAAGATGCAGAGCTATATGACCA | CTGACCTTCACGCAACAGAC |
| Ego221 | PPIP5K2 | 17,622,251 | 17,623,325 | CTTATGCAAAACTCAGGAATCCA | CGGTTCAGAACTGCATAGCG |
| Ego222 | NUDT12 | 17,750,608 | 17,751,643 | TGACTTGCATTATCACAACAGGA | TTGTCCCAGTCTCCAAGGAA |
| Ego225 | SNX2 | 23,301,475 | 23,302,587 | AAAGTGACCTGCGCATCTTG | CATTCGTTTGATTGCTGCAGT |
| Ego226 | RASA1 | 24,563,908 | 24,565,046 | ACTGGGAGTGTATGAGCTTCT | ACCAGTCCAGGAACGTCAAA |
| Ego228 | TRIM36 | 27,126,184 | 27,127,253 | CAACCATGAGCACTGCCTAC | CACCTGGCTCTCCTTACTGA |
| Ego231 | TSTD2 | 31,655,184 | 31,656,169 | TACTGCACAGGAGGGATTCG | GGTAAACCTCTCTGCACACTG |
| Ego233 | PIK3C3 | 34,847,680 | 34,848,678 | AGCGCATAACTCTGACAGACT | CAGAGGGACCCAAAGACTCA |
| Ego236 | MTMR12 | 40,467,025 | 40,468,074 | CACAGCACAGATCAGTAGCAC | AGAAGTTGCTGAGTGTCTGGA |
| Ego237 | LIFR | 42,670,413 | 42,671,542 | GCGACAGGGATGAGAATTGC | GCCCTCCAAACAGCCTCTAT |
| Ego240 | DDX4 | 47,480,469 | 47,481,564 | CCAGGAAGGCTTCTAGACATCA | CCCATATCAAGCATGCGGTC |
| Ego241 | ERCC8 | 49,383,661 | 49,384,825 | AGCAGGTGAAGTCCATCATTTG | AGAGCTTCTGGATGTCTGAGT |
| Ego242 | CWC27 | 50,580,730 | 50,581,714 | AAGTCCAAAGTAAAAGGCACAAA | CCAAAGGAAAGCAGGTTAAAGT |
| Ego243 | RASEF | 53,009,943 | 53,011,046 | CTGAAGCGCCAGTATGACAC | CAGTTCAACTACATCTTCCCTGT |
| Ego246 | FOCAD | 57,707,626 | 57,708,729 | GTGTCACAGCCAGTTCAAGG | AGCCATCAGTATCCACCTGT |
| Ego247 | SNAPC3 | 59,927,625 | 59,928,726 | TCCTATCCAAACAGTCATCATGA | ACCTTTACTGTCACCAAGGGA |
| Ego248 | similar to MPDZ | 60,830,550 | 60,831,630 | GCAGGACTAACTTCAACTGGC | TGGTCTTGTACTGAGGAGGT |
| Ego250 | CDC37L1 | 64,135,894 | 64,136,869 | AGATCTCCTGTGTTCTGCAAC | CTCAGTGTCAGTCCTGGTGT |
| Ego251 | DOCK8 | 65,603,665 | 65,604,686 | CAACCACTGACTCAAAGGCA | GCTGGCTGTACAACCTATGG |
| Ego255 | DHFR | 72,301,146 | 72,302,156 | TCCTGAGAAGAACCGTCCTT | AGATAATGTGCTCCTTTCGGG |

**Supplementary Table 9 |** Point estimates of polymorphism statistics for the *Red* locus, and mean values at the reference loci. $\pi$, nucleotide diversity; $D_{Taj}$, Tajima's *D*; $F_{ST}$, differentiation between the red and black allelic classes; $d_{XY}$, mean pairwise diversity between the red and black allelic classes. Figures in brackets denote the standard error, where applicable.

| | | | Within Allelic Class | | | Classes Combined | | Between Classes | |
|---|---|---|---|---|---|---|---|---|---|
| Region | No. of loci | (mean) locus length (bp) | Class | $\pi$ | $D_{Taj}$ | $\pi$ | $D_{Taj}$ | $F_{ST}$ | $d_{XY}$ |
| Reference loci | 24 | 1,041.5 | red | 0.0032 (0.0005) | -0.304 (0.1234) | 0.0033 (0.0004) | -0.818 (0.1270) | 0.038 (0.0227) | 0.0033 (0.0004) |
| | | | black | 0.0033 (0.0005) | -0.599 (0.1503) | | | | |
| *Red* locus | 1 | 72,000 | red | 0.0004 | -0.844 | 0.0102 | 2.193 | 0.951 | 0.019 |
| | | | black | 0.0015 | -0.891 | | | | |

**Supplementary Table 10 |** Details of the nine microsatellite loci used to test for global differentiation among individuals in the Wyndham population sampled in 2008. Included are the locus name, sample size (*N*), number of alleles (*k*), significance of a test for HWE (NS not significant, ND not done), chromosome (Chr), observed allele size ranges and primer sequences.

| Locus | *N* | *k* | HWE | Chr | Allele size range (bp) | Primer sequences and labels (5' → 3') |
|---|---|---|---|---|---|---|
| TG01-000 | 160 | 9 | NS | 1A | 251-267 | [6-FAM]TTGCTACCARAATGGAATGT<br>TCCTAACCATGAGAAGCAGA |
| TG01-147 | 161 | 9 | NS | 1 | 273-291 | [5HEX]TGAGCCACTACAGAGTGGAAA<br>GCCACTACAATGAAGAAAATATTACAG |
| TG02-078 | 161 | 6 | NS | 2 | 290-300 | [5HEX]TGTTAAAGCCTGTTCCATAGG<br>TTCCCCATAAAGTATGTACGC |
| TG03-031 | 160 | 7 | NS | 3 | 199-215 | [6-FAM]ATTGCACATGAACCTGGAAG<br>TCATTACTTGAAGCAGGTCTCTG |
| TG04-012 | 158 | 14 | NS | 4A | 119-168 | [5HEX]TGAATTTAGATCCTCTGTTCTAGTGTC<br>TTACATGTTTACGGTATTTCTCTGG |
| Ase24-ZFS | 161 | 8 | NS | 5 | 199-216 | TGTGCATGTGTGTGCATTTG<br>TGTTGTCTGAAGCTGTCATTGC |
| Tgu07 | 161 | 8 | NS | 6 | 99-113 | [5HEX]CTTCCTGCTATAAGGCACAGG<br>AAGTGATCACATTTATTTGAATAT |
| Tgu02 | 159 | 3 | NS | 7 | 184-188 | [5HEX]TGGATTACCTGTCTGAAAGACC<br>TTCACTGTCTAGTCCAACCCTGT |
| TG22-001 | 157 | 29 | ND | 22 | 255-286 | [5HEX]TTGGATTTCAGAACATGTAGC<br>TCTGATGCAAGCAAACAA |

**Supplementary Table 11 |** $L_n$-likelihood results and parameter estimates from the method of Zeng and Charlesworth[24] to test for a change in population size, and for non-neutral evolution. Parameters: $u$ – the population-scaled AT → GC mutation rate; $v$ – the population-scaled GC → AT mutation rate; $\gamma$ – the population-scaled selection coefficient favouring AT alleles; $a$ – the ratio of population sizes $N_1/N_0$, where $N_0$ is the population size before the change and $N_1$ is the population size after the change, forward in time; $\tau$ – the time in units of $2N_0$ generations into the past at which the inferred change in population size occurred.

| Model | Max ln-likelihood | -2Δ lnL[a] | $u$ | $v$ | $\gamma$ | $a$ | $\tau$ |
|---|---|---|---|---|---|---|---|
| Equilibrium | -1.6729e+04 | | 1.68e-03 | 6.51e-03 | -7.56e-01 | n/a | n/a |
| Size change 1 step, $\gamma = 0$ | -1.6713e+04 | 30.6870 | 1.31e-03 | 2.43e-03 | n/a | 6.91e+00 | 9.99e-02 |
| Size change 1 step | -1.6708e+04 | 9.7259 | 1.00e-03 | 3.38e-03 | -6.11e-01 | 6.58e+00 | 9.51e-02 |

[a] Two times the difference in log-likelihoods between this model and the next-worst model (the model above in the table).

**Supplementary Table 12 |** Testing $R_M$ with known $\rho$.

| $\rho$ ($bp^{-1}$) | Replicate | $R_M$ | |
|---|---|---|---|
| | | DAF = 0.1 | DAF = 0.9 |
| 0 | 1 | 0 | 0 |
| | 2 | 0 | 0 |
| | 3 | 0 | 0 |
| | 4 | 0 | 0 |
| | 5 | 0 | 0 |
| | 6 | 0 | 0 |
| | 7 | 0 | 0 |
| | 8 | 0 | 0 |
| | 9 | 0 | 0 |
| | 10 | 0 | 0 |
| 0.0002 | 1 | 5 | 6 |
| | 2 | 3 | 12 |
| | 3 | 5 | 10 |
| | 4 | 7 | 9 |
| | 5 | 8 | 4 |
| | 6 | 14 | 4 |
| | 7 | 9 | 5 |
| | 8 | 2 | 7 |
| | 9 | 9 | 5 |
| | 10 | 8 | 10 |
| 0.002 | 1 | 34 | 42 |
| | 2 | 48 | 38 |
| | 3 | 36 | 51 |
| | 4 | 37 | 49 |
| | 5 | 41 | 36 |
| | 6 | 38 | 37 |
| | 7 | 41 | 38 |
| | 8 | 44 | 46 |
| | 9 | 38 | 38 |
| | 10 | 35 | 32 |

**Supplementary Table 13 |** Recombination in the region encompassing the *Red* locus.

| Samples[§] | both | | black | | red | |
|---|---|---|---|---|---|---|
| Locus[†] | 100-kbp | *Red* | 100-kbp | *Red* | 100-kbp | *Red* |
| $R_M$[‡] | 58 | 41 | 21 | 15 | 8 | 4 |
| $R_M$ bp$^{-1}$ | 5.82 x $10^{-4}$ | 5.69 x $10^{-4}$ | 2.11 x $10^{-4}$ | 2.01 x $10^{-4}$ | 0.8 x $10^{-4}$ | 0.55 x $10^{-4}$ |

[§], The alleles included in the assay. both: all 12 sequences, six each of the red and black alleles; black: six black alleles; red: six red alleles. [†], The region on which the test was performed. 100-kbp: the entire 99,669 bp region for which we had sequence data; *Red*: the 72,000-bp region within the 100 kbp that corresponds to positions 12,501–84,500 bp of the sequenced region where $F_{ST} > 0.8$ between the red and black alleles. [‡], The $R_M$ statistic of Hudson and Kaplan[25].

**Supplementary Table 14 |** Recombination statistics for 24 Z-linked intronic reference loci. Values of $\rho$ were obtained using LDhat. Values of $R_M$ were obtained using RecMin. *Length of sequence after trimming primer sequences used in analysis.

| Loci | Target | Length* (bp) | $\rho$ ($bp^{-1}$) | $R_M$ ($locus^{-1}$) |
|---|---|---|---|---|
| Ego210 | *KATNAL2* | 1,076 | 0.0195 | 2 |
| Ego213 | *LPL* | 1,156 | 0.0156 | 3 |
| Ego214 | *ADAMTS19* | 965 | 0.0715 | 5 |
| Ego216 | *CNTNAP4* | 1,075 | 0.0437 | 2 |
| Ego217 | *DAPK1* | 972 | 0.0309 | 0 |
| Ego221 | *PPIP5K2* | 1,043 | 0 | 0 |
| Ego222 | *NUDT12* | 970 | 0.0505 | 0 |
| Ego225 | *SNX2* | 1,078 | 0 | 0 |
| Ego226 | *RASA1* | 1,160 | 0.0983 | 0 |
| Ego228 | *TRIM36* | 1,056 | 0 | 0 |
| Ego231 | *TSTD2* | 952 | 0.0032 | 1 |
| Ego233 | *PIK3C3* | 953 | 0.0021 | 0 |
| Ego236 | *MTMR12* | 995 | 0.002 | 0 |
| Ego237 | *LIFR* | 1,090 | 0 | 0 |
| Ego240 | *DDX4* | 1,050 | 0.0057 | 1 |
| Ego241 | *ERCC8* | 1,123 | 0.0018 | 0 |
| Ego242 | *CWC27* | 931 | 0.0054 | 1 |
| Ego243 | *RASEF* | 1,049 | 0.001 | 0 |
| Ego246 | *FOCAD* | 1,055 | 0 | 0 |
| Ego247 | *SNAPC3* | 1,054 | 0 | 0 |
| Ego248 | *similar to MPDZ* | 1,033 | 0 | 0 |
| Ego250 | *CDC37L1* | 880 | 0.0273 | 0 |
| Ego251 | *DOCK8* | 866 | 0.0231 | 0 |
| Ego255 | *DHFR* | 978 | 0.0409 | 2 |

**Supplementary Methods**

**Test for neutrality at the *Red* locus** In this work, we aim to investigate whether or not the patterns of genetic diversity observed within and between head colour morphs at the *Red* locus in the Gouldian finch are the result of natural selection. It is inappropriate to apply standard population genetic approaches that assume random sampling to non-random samples, as in our case where the red and black alleles are each represented in our sample sequenced on the MiSeq at 50%, which is not consistent with the allele frequency in natural population (see below). In order for us to have included the rare red allele at natural allele frequencies, an impractically large sample size would have been required. To overcome the problems introduced by our non-random sample, we employ coalescent simulations to generate patterns of polymorphism under the neutral expectation, conditioning on our sampling scheme and the allele frequencies in the wild population at the *Red* locus. We estimate population genetic parameters including the population mutation rate ($\theta = 4N_e u$) and the population recombination rate ($\rho = 4N_e r$) across the ~100-kbp sequenced region, taking into account demography and other non-selective forces, which we infer from Z-linked intronic loci unlinked to the *Red* locus. It is important to consider demography when testing for selection, because the effect of demographic events on genetic polymorphism may be mistaken for the effect of selection[26-28]. Similarly, GC-biased gene conversion (gBGC) has the potential to distort the site frequency spectrum (SFS) in such a way that may lead to incorrect inferences of selection and/or demography[29-31]. gBGC has been suggested to play a part in the evolution of bird genomes[32-34], so we attempt to quantify its effect in this study at the same time as making demographic inferences. We also implement a modified version of the HKA test[35] that accounts for non-random sampling and recombination at one locus, as well as uneven lengths between loci.

**Testing for a change in population size and gBGC** We used the matrix method of Zeng and Charlesworth[24] (see also Evans *et al*.[36]) to estimate the likelihood of a one-step change in population size, based on our 24 Z-linked intronic reference loci. Briefly, this method uses the site-frequency spectrum to infer parameters of a two-allele model with reversible mutation, selection and/or gBGC, and changes in population size. We defined A/T and G/C as our two alleles. We define $u$ as the rate at which A/T alleles mutate to G/C alleles, $v$ as the mutation rate in the opposite direction, and $\kappa = v/u$ as the mutation bias parameter. The selection coefficient, $\gamma$, is defined as $\gamma = 4N_e s$ where $N_e$ is the effective population size and $s$ is the selection coefficient against heterozygous carriers of the G/C allele. To model a change in population size, we assume that the population in the past is at equilibrium with population size $N_1$, which then changes instantly to $N_0$ (this can be either an increase or a reduction in size) and remains in this state for $t$ generations until a sample is taken from the population in the present day[24,36,37]. For each model, in order to ensure that the true MLE was found, we ran the search algorithm multiple times (typically 1,000), each initialised from a random starting point. All the reported results were found by multiple searches with different starting conditions.

The one-step population size change model fitted the data significantly better than the equilibrium model, given the data from our reference loci ($\chi^2 = 40.41$, d.f. = 2, $P < 0.001$; Supplementary Table 11). The time of the size change, $t$, was estimated as 9.51e-2 × $2N_0$ generations into the past, and the population was inferred to have undergone an expansion (forward in time) at this point by a factor of $a$ = 6.58 (Supplementary Table 11). Including a second size change did not result in a significant increase in the log-likelihood of the model over a single size change ($\chi^2 = 1.12$, d.f. = 2, $P = 0.57$; full data not shown). The one-step size change model was also a significant improvement on a one-step size change model with $\gamma = 0$ ($\chi^2 = 9.73$, d.f. = 1, $P = 0.002$; Supplementary Table 11) i.e., a model incorporating selection (or some other force that favours one of our two allelic classes, A/T vs. G/C) fits the data better. The inferred value of $\gamma$ was -0.611, which indicates a force favouring G/C alleles over A/T alleles. The mutation bias parameter, $\kappa = \frac{GC \rightarrow AT}{AT \rightarrow GC} = 3.37$ under the best-fit model. This result is close to the value reported in the *Ficedula* flycatcher lineage by Bolívar *et al*. of 3.41[34]. Because the sequences we used to conduct these tests were intronic, the significant value for $\gamma$ suggests the action of GC-biased gene conversion (gBGC) as opposed to, for example, selection for preferred codons. gBGC has been more thoroughly investigated in mammals than in birds, and its underlying mechanisms are best characterised in yeast[30], but its effect on avian genome evolution may be important[34]. gBGC was not incorporated into the coalescent simulations described below due to limitations of the methods we used.

**Maximum-likelihood estimates of allele frequencies** Testing for departures from neutrality at the *Red* locus requires information about frequencies of the $Z^r$ and $Z^R$ alleles (for brevity, hereafter subscripts '$r$' and '$R$' refer to the $Z^r$ and $Z^R$ alleles, respectively) at the *Red* locus, denoted as $p_r$ and $p_R$, respectively. Wild birds ($n$ = 161) genotyped at the *Red* locus did not show evidence of departure from the Hardy–Weinberg Equilibrium (Supplementary Table 3). We calculated the ln-likelihood of $p_r$ and $p_R$ as

$$L(D|p_r, p_R) = 2X_{rr} ln\,(p_r) + X_{rR} ln\,[2p_r p_R] + 2X_{RR} ln\,(p_R) + Y_r ln\,(p_r) + Y_R ln\,(p_R) \qquad (1)$$

where $D$ represents the genotypes of the 161 birds, $X_{rr}$ is the observed count of males homozygous for $Z^r$, $X_{rR}$ is the observed count of heterozygotes in males, $X_{RR}$ is the observed count of males homozygous for $R$, $Y_r$ is the observed count of females hemizygous for $Z^r$ and $Y_R$ is the observed count of females hemizygous for $R$. Subject to maximising Supplementary Equation 1, with the constraints $p_r + p_R = 1$ and $0 \leq p_r \leq 1$, our maximum-likelihood estimates of the allele frequencies for the $Z^r$ and $Z^R$ alleles were 0.856 (95% CIs: 0.808, 0.897) and 0.144 (95% CIs: 0.103, 0.192), respectively.

**Estimating $\theta$ under a change in population size** We sought to obtain an estimate of $\theta$, which is defined as $4N_e u$, where $N_e$ is the effective population size and $u$ is the mutation rate per site, under an inferred change in population size obtained from the method of Zeng and Charlesworth[24] (see above). To do so, we wanted to determine a value of $\theta$ such that the expected level of diversity under this population-scaled mutation rate, given the inferred one-step change in population size, equals the mean value of $\pi_{\text{total}}$ observed at our 24 reference loci (Supplementary Table 9).

The expected value of diversity at a neutral site is $\theta E(X)$, where $X$ is the time in units of $2N_0$ generations, and $N_0$ is the population size after the change, to the most recent common ancestor for two randomly chosen alleles taken from the current population. If an expression of $\theta E(X)$ can be found, we can estimate $\theta$ using

$$\hat{\theta} = \frac{\pi_{obs}}{E(X)} \tag{2}$$

The coalescent rate in $[0, \tau]$ is 1 and that in $[\tau, \infty]$ is $a = N_0/N_1$, where $\tau$ is the time in units of $2N_0$ generations between the time of the inferred change in population size and the present (when a sample was taken from the population), $N_0$ is the current population size and $N_1$ is the population size before the inferred change in population size. Furthermore, the probability of coalescing in $[\tau, \infty]$, referred to as $c$, can be calculated as

$$c = \int_{\tau}^{\infty} e^{-t} dt = e^{-\tau} \tag{3}$$

Thus, the probability density function, $f(t)$, for the coalescent time $X$ is

$$f(t) = \begin{cases} e^{-t}, & \text{if } 0 \leq t \leq \tau; \\ cae^{-a(t-\tau)}, & \text{if } t > \tau. \end{cases} \tag{4}$$

Thus, $E(X)$ can be calculated as

$$\begin{aligned} E(X) &= \int_{0}^{\tau} te^{-t} dt + \int_{\tau}^{\infty} tcae^{-a(t-\tau)} dt \\ &= 1 - e^{-\tau}(1+\tau) + \frac{e^{-\tau}(1+a\tau)}{a} \end{aligned} \tag{5}$$

Replacing $\tau$ and $a$ in Supplementary Equation 5 with the MLEs from the Zeng and Charlesworth[24] method, we obtain $E(X)$, which is then inserted into Supplementary Equation 2 to obtain $\hat{\theta}$. Using the maximum-likelihood estimates of $\tau$ (the time to the size change in units of $2N_0$ generations; see above) and $a$ (the factor by which the population increased) we can obtain $E(X) = 0.2362$ from Supplementary Equation 5. Since $\pi_{obs} = 0.0033$ (Supplementary Table 9), from Supplementary Equation 2 we have $\hat{\theta} = 0.0144$.

**Estimating $\rho$, the population recombination rate** In the following analyses we used the ~100-kbp sequenced region that included the *Red* locus. We did not delimit the locus more tightly to the Red locus because:

i) As shown clearly in Figure 3, the evidence for selection on variants within the red locus comes from the observed patterns of variability (black lines) lying well outside what is expected under neutrality (grey area). If we restricted the simulation to the 72-kbp *Red* locus then these patterns would not be apparent.

ii) In this context, the fact that the patterns of variability in the flanking regions fall well within what is expected under neutrality highlights that what we see at the *Red* locus falls outside of neutral expectation, and also adds credibility to our simulation model.

iii) The presence of these flanks also enables a visual appreciation of the extent of the *Red* locus and avoids the need to make a slightly arbitrary decision about the precise extent of the *Red* locus.

We inferred the population recombination rate, $\rho$ ($4N_e r$, where $N_e$ is the effective population size, and $Z^r$ is the recombination rate per site), using polymorphism. For the reference loci, LDhat[38] or LDhelmet[39] were employed. Because our sample at the *Red* locus is not a random representation of the population, we refrained from obtaining estimates of $\rho$ at the *Red* locus using these methods. To get around this difficulty, we estimated the $R_M$ statistic of Hudson and Kaplan[25], which tests for the number of recombination events that can be parsimoniously inferred from a sample of DNA sequences, using the computer package *RecMin*[40] (The $R_h$ statistic of Myers and Griffiths gave essentially the same results; data not shown). Importantly, provided that there has been no recurrent mutation, $R_M$ is non-zero only when there have been recombination events in the history of the sample[40] (Supplementary Table 12). Thus, $R_M$ should be a robust test for the presence of recombination.

The following procedure was used to obtain $\rho$ values that were likely to be compatible with data for the sequenced region. We first simulated a neutral locus equivalent to the sequenced region using mbs[41], conditioning on our sampling scheme (six each of $Z^r$ and $Z^R$ alleles, from the wild allele frequency distribution of 0.856 and 0.144, with each allele being treated as derived in turn). To generate neutral variants, the simulations were conducted using the estimate of $\theta$ from our 24 reference loci under the size change model (see the previous section), and incorporating the inferred demographic expansion (see above). A range of values of $\rho$, from 0 to 0.005 $bp^{-1}$, was considered. We performed 1,000 simulations for each combination of parameters. We then estimated the $R_M$ on each simulated dataset and obtained the mean across replicates. The mean values of $R_M$ were plotted against $\rho$, and $\rho$ values with average $R_M$ comparable to the observed $R_M$ were regarded as compatible with the data.

Using the procedure above, we estimated the minimum parsimonious number of recombination events ($R_M$) in the entire sequenced region as 58 (Supplementary Table 13), from which we inferred $\rho \approx 0.004$, including the inferred demographic expansion, where $\rho = 4N_0r$ and $N_0$ is the present-day population size (see above; Supplementary Figure 8). The inferred value of $\rho$ was robust to the sections of the sequenced region for which we calculated the observed $R_M$. The $R_M$ $bp^{-1}$ does not differ between the *Red* locus and its flanking regions (goodness of fit $\chi^2_1 = 0.05$, $P = 0.82$; Supplementary Table 13). We assumed a single value of $\rho = 0.004$ across the sequenced region for all analyses incorporating the expansion below. This estimate lies within the estimates of $\rho$ $bp^{-1}$ at the reference loci (Supplementary Table 14). The value of $\rho$ was also comparable ($\rho = 0.001$) when a constant-size population was simulated (data not shown).

**Coalescent simulations of the sequenced region – under a population size change** In order to assess whether the patterns of genetic polymorphism at the *Red* locus are compatible with purely neutral processes we conducted simulations using mbs[41]. We simulated a 99,669-bp region (equivalent in length to the sequenced region), using the estimate of $\theta$ from our 24 reference loci under the size change model (Supplementary Table 11) and the estimate of $\rho$ = 0.004 from the Zeng and Charlesworth[24] method described in the section above. We replicated the missing data in our observed dataset by masking the same sites in our simulated datasets before calculating summary statistics. We carried out two sets of simulations, treating each allele (*r* and *R*) as derived, in turn.

As a first test, we examined whether patterns of diversity at the *Red* locus, as summarised by sliding-window plots of $F_{ST}$, $D_{XY}$, $\pi$ and Tajima's *D*, could be explained by a neutral model with a non-random sampling scheme and the inferred demographic expansion, as well as the inferred values of $\theta$ and $\rho$. Assuming a DAF of 0.144 at the focal site (the $Z^R$ allele is derived), the observed data for the sequenced region lie above the 95% confidence intervals (CIs) of our coalescent simulations for many windows, for a range of summary statistics that incorporate both allelic classes (Fig. 3a-d). For the statistics calculated using within-allelic classes ($D_{\text{Taj red}}$, $D_{\text{Taj black}}$, $\pi_{\text{red}}$, $\pi_{\text{black}}$), in most cases, the observed data generally fall within the 95% CIs of the simulated distributions (Supplementary Figure 9), probably in part caused by their higher variance because they were calculated on smaller sample sizes. The exception was $\pi_{\text{red}}$, which lies below the simulated 95% CIs of $\pi_{\text{red}}$ (Supplementary Figure 9b). Balancing selection is expected to lead to reduced diversity at neutral sites linked to the site under selection within the less frequent allelic class, but on a much smaller scale relative to the increase in polymorphism when allelic classes are combined[42,43]. These predictions do not account for demographic changes, however, such as the one we suggest has occurred in the Gouldian finch, and consequently it is unclear to us whether the observed level of $\pi_{\text{red}}$ is compatible with the action of balancing selection alone. These results are qualitatively identical for a DAF of 0.856 (Supplementary Figure 10). Overall, these simulations suggest that there are multiple aspects of the data that cannot be explained by the sampling scheme alone.

**A modified HKA test incorporating non-random sampling and recombination** We sought to modify the HKA test[35] for four reasons: (1) our sample of alleles at the *Red* locus is non-random, which is not allowed for in the original formulation of the test; (2) *P*-values from the $\chi^2$ distribution are not robust when we have unequal locus sizes (see Supplementary Figure 15); (3) simulation-based *P*-values assuming no recombination in the *Red* locus may be too conservative, given its size and clear evidence of intra-locus recombination; and (4) a sliding-window approach may help to identify candidate sites by contrasting the results of the test for different regions of the sequenced region, whilst retaining the same reference loci as a comparison.

We also carried out simulations to assess the effect of a one-step change in population size on the HKA test, and found that, in the presence of a demographic expansion equivalent to that which we inferred from our data, not directly accounting for the size change renders the HKA test conservative (Supplementary Figure 16). Therefore we chose not to incorporate the change in population size in our HKA analysis.

We first set out to estimate the expectation and the variance of the number of segregating sites within a window of the sequenced region, given the sampling scheme and the location of the focal site. Assuming the infinite-sites model and defining $\theta$ *per window* as $4N_e u$, the mean and variance of the number of segregating sites in this window, *S*, are given by

$$E(S) = \theta E(T) \tag{6}$$

and

$$Var(S) = E[Var(S|T)] + Var[E(S|T)] = \theta E(T) + \theta^2 Var(T) \tag{7}$$

where $T$ is the sum of the total branch length across all nucleotide sites in the window. The above equations have made use of the fact that, given $T$, $S$ is Poisson-distributed with mean $\theta T$.

In order to obtain $E(T)$ and $Var(T)$ for our sampling scheme and the recombination rate at (a window of) the sequenced region, we used the following algorithm:

1) Set $\theta_{\text{sim}}$ *per window* to 1.0 (the choice of $\theta_{\text{sim}}$ is unimportant)
2) For $i$ from 1 to 1,000,000

   Do

   a) Use SelSim and $\theta_{\text{sim}}$ to generate a random sample conforming to the sequenced region (in terms of derived allele frequency, sample size at each allele class, recombination rate, length and the site at which sampling is conditioned upon in the centre of the locus)
   b) Record the number of segregating sites in each window of the sequenced region and store it in $X_i$

   Done

To conduct SelSim simulations of the sequenced region we simulated a 99,669-bp region, placing the focal site in its centre (although this choice does not materially affect the results), and using the same sampling scheme as in the observed data (six each of the two allele classes). We set $\rho$ = 0.001 (see Estimating $\rho$, the population recombination rate), the value of $\theta$ (*per window*) as 1 and used point estimates of the two DAFs (0.144 and 0.856). As with the position of the focal site, our results were robust to this value of $\rho$. We did not sample DAFs from their posterior distributions primarily because it was computationally intractable to perform the large number of simulations needed to estimate $E(T)$ and $Var(T)$. Furthermore, the estimates of DAF have fairly small 95% confidence intervals (see Maximum-likelihood estimates of allele frequencies).

Using the number of segregating sites ($X_i$s) obtained from the simulations described above, we obtained estimates of $E(T)$ and $Var(T)$, denoted as $\mu_T$ and $\sigma_T^2$, respectively, as follows

$$\bar{X} = \frac{1}{n}\sum_{i=1}^{n} X_i \tag{8}$$

where $n$ is the number of simulations performed in total

$$\mu_T = \bar{X}/\theta_{sim} \tag{9}$$

$$Y^2 = \frac{1}{n-1}\sum_{i=1}^{n}(X_i - \bar{X})^2 \tag{10}$$

$$\sigma_T^2 = \frac{Y^2 - \theta_{sim}\mu_T}{\theta_{sim}^2} \tag{11}$$

Since $\mu_T$ and $\sigma_T^2$ were treated as known quantities in the analysis below, to minimise statistical noise, we estimated them using $10^6$ simulations.

Now equations (5) of Hudson *et al.*[35] for the case when we have polymorphism data from only one species can be rewritten as

$$\sum_{i=1}^{M-1} s_i + s_R = \sum_{i=1}^{M-1} C(n_i)\hat{\theta}_i + \hat{\theta}_R\mu_T, \tag{12}$$

$$\sum_{i=1}^{M} D_i = \left(\hat{Y}+1\right)\sum_{i=1}^{M}\hat{\theta}_i,$$

$$s_i + D_i = \hat{\theta}_i\left\{\hat{Y} + 1 + C(n_i)\right\}, \qquad i = 1, \dots, M-1,$$

where $M$ is the total number of loci, among which the first $M$ - 1 are our reference loci, and the $M^{\text{th}}$ locus is the focal window of the sequenced region, $C(n_i)$ is $\sum_{j=1}^{n_i-1} 1/j$ where $n$ is the sample size at the $i^{\text{th}}$ locus, $D_i$ is the observed divergence between the ingroup data and the outgroup data, and $Y$ is the divergence time between the ingroup and outgroup in units of $2N$ generations. In Supplementary Equation 12, hats have been added to parameters that are to be estimated, whereas all other symbols represent values obtained from the sequence data.

We carried out the test above as a sliding-window across the sequenced region, using a window size of 1,000 bp and a step size of 500 bp. For each given window, we solved the $M$ + 1 unknowns in Supplementary Equation 12 using a custom script in R, and calculated the $X^2$ statistic as in Hudson *et al.*[35], except that we obtained the mean and variance of tree lengths ($T$) from Supplementary Equations 9 and 11, above, then used our estimate of $\theta$ along with Supplementary Equations 6 and 7 to obtain the mean and variance of $S$.

In order to test the significance of $X^2$ we used two methods, (i) the chi-squared distribution with $M$ – 1 degrees of freedom and (ii) by conducting coalescent simulations. The procedure for conducting coalescent simulations for the reference loci is similar to that described in Hudson *et al.*[35]. The main difference is that, for a window of the sequenced region, we simulated polymorphism data using the algorithm given above, but using our estimate of $\theta$ in place of $\theta_{\text{sim}}$. We simulated divergence at each locus ($D_i$) as

$$D_i = Poisson(\hat{\theta}_i \hat{Y}) + Geometric\left(\frac{1}{1+\hat{\theta}_i}\right) \tag{13}$$

where $\hat{\theta}_i$ and $\hat{Y}$ were estimates obtained by solving Supplementary Equation 12 above[44]. To estimate the $P$-value, we carried out 1,000 simulations for each window and estimated the probability that the value of $X^2$ from the simulation procedure was greater than or equal to our observed value of $X^2$. We carried out the above procedure twice, once each for a DAF of 0.144 and of 0.856 at the focal site.

For the 2,781-bp portion of the *Red* locus for which we do not have divergence data, which coincides with a transposable element (TE) insertion specific to the Gouldian finch lineage (and thus not present in the zebra finch outgroup), we estimated divergence using 3 kbp (1.5 kbp from each side) of the *Red* locus immediately adjacent to the missing data. This procedure returned a value for divergence of 52.1 $kbp^{-1}$, which is greater than the mean value of the windows of the sequenced region for which outgroup sequence data are available (46.1 $kbp^{-1}$), making our HKA test conservative for this region in comparison to using a locus-wide average value of divergence.

Analysis of the sequenced region based on $R_M$ gave clear evidence of recombination. By carrying out a sliding-window implementation of the test it is possible to contrast the results of the test between different regions of the sequenced region whilst retaining the same reference loci as a comparison. In this way, we were able to identify regions showing the most unusual patterns of polymorphism.

Our sliding-window implementation of the HKA test, taking into account non-random sampling and recombination at the sequenced region, reveals significant departures from neutrality for multiple windows of the sequenced region (Fig. 3e). *P*-values for both DAFs of 0.144 and 0.856, and both the chi-squared and the simulation methods, are all in close accord. The most significantly departed region coincides with the region of the *Red* locus for which we do not have divergence data from zebra finch due to the insertion of a putative TE in the Gouldian finch lineage, and at which the levels of polymorphism within the Gouldian finch lineage are highest (Fig. 2b). However, we obtain similar results elsewhere in the *Red* locus if we treat the TE region as the single focal site (by removing polymorphism data from that region) (Supplementary Figure 17).
